# PIP_2_ stabilizes Na_V_1.5 gating and links receptor signaling to cardiac late sodium current

**DOI:** 10.64898/2026.06.29.735321

**Authors:** Kirin D. Gada, Jordie M. Kamuene, Ana Santa Cruz, Ziyue Meng, Jenna G. Connolly, Florence Ng, Xinyi Ma, Aishwarya Chandrashekar, Yu Xu, Meng Cui, Leigh D. Plant

## Abstract

The cardiac sodium channel Na_V_1.5 initiates each heartbeat by generating the rapid depolarizing upstroke of the action potential. Dysregulation of Na_V_1.5 gating can produce cardiac arrhythmias by slowing inactivation, increasing late sodium current (I_Na,L_), and impairing electrical stability. Here, we show that phosphatidylinositol-4,5-bisphosphate (PIP_2_) is a critical membrane cofactor that stabilizes Na_V_1.5 gating. Acute PIP_2_ depletion in human iPSC-derived cardiomyocytes, produced by activation of endogenous AT1 receptors, activation of an engineered M3q-DREADD, or optogenetic recruitment of CRY2-pseudojanin, shifted voltage dependence, slowed fast inactivation, and increased I_Na,L_. These effects were prevented by augmenting intracellular PIP_2_, required PLC activity when driven by Gq-coupled receptors, and were independent of downstream Ca² or PKC signaling. Unlike the skeletal-muscle isoform Na_V_1.4, Na_V_1.5 displayed PIP_2_-dependent shifts in both activation and steady-state inactivation, indicating isoform-specific lipid coupling. Induced-fit docking and molecular dynamics simulations identified a PIP_2_-interaction interface between the domain IV voltage sensor and pore that contains disease-linked residues. The disease-reported variant R1644C weakened and redistributed the predicted PIP_2_-contact network, produced elevated basal I_Na,L_, showed enhanced sensitivity to PIP_2_ depletion, and caused an approximately 30-fold reduction in apparent functional PIP_2_ sensitivity in excised patches. These findings define a lipid-dependent mechanism that stabilizes Na_V_1.5 gating and reveal how physiological Gq signaling and inherited channel variants can converge on the channel-PIP_2_ axis to promote proarrhythmic late sodium current.

## Introduction

Electrical activity in the human heart is initiated by activation of the voltage-gated sodium channel Na_V_1.5, which drives the rapid depolarizing upstroke of the cardiac action potential (1). The coordinated opening and inactivation of Na_V_1.5 determine conduction velocity and contribute to repolarization reserve, and even small disturbances in channel gating can predispose to arrhythmia (2, 3). In addition to the transient inward sodium current (I_Na_), Na_V_1.5 generates a much smaller late current (I_Na,L_) that persists during the plateau phase of the action potential. Elevation of I_Na,L_, whether through inherited variants, metabolic stress, or pathological signaling, prolongs the action potential and contributes to ventricular and atrial arrhythmias (4, 5). Mechanisms that stabilize Na_V_1.5 gating and restrain I_Na,L_ are therefore central to the electrical stability of the myocardium (6, 7).

Phosphatidylinositol-4,5-bisphosphate (PIP_2_) is a minor yet essential component of the inner leaflet of the plasma membrane that acts as a cofactor for many ion channels (8). Inward-rectifier and KCNQ potassium channels require PIP_2_ to remain open, and loss of membrane PIP_2_ through phospholipase C (PLC) activity leads to rapid current rundown (9). Despite the ubiquity of PIP_2_ regulation in ion-channel physiology, its role in cardiac sodium channels has remained comparatively unexplored. We recently showed that targeted dephosphorylation of PIP_2_ in the skeletal muscle sodium channel Na_V_1.4 shifts the voltage dependence of activation to more negative potentials and increases I_Na,L_ without detectably altering steady-state inactivation, consistent with lipid-dependent stabilization of Na_V_ gating (10). These findings raised the broader hypothesis that PIP_2_ modulates voltage-gated sodium channel function, but whether this mechanism applies to the closely related cardiac isoform Na_V_1.5 in cardiomyocytes, and whether it has pathophysiological relevance, remained unknown.

In cardiomyocytes, PIP_2_ levels are dynamically controlled by Gq-coupled signaling. Activation of Gq-coupled receptors engages PLC to hydrolyze PIP_2_ into diacylglycerol (DAG) and inositol-1,4,5-trisphosphate (IP_3_), transiently reducing the membrane pool of the lipid. The relative contribution of direct PIP_2_ loss versus secondary signaling through DAG, PKC (Protein Kinase C), or Ca² , however, has been difficult to isolate (11–13). Here, we use human iPSC-derived cardiomyocytes and heterologous expression systems to dissect these contributions by combining optogenetic phosphatase control, chemogenetic activation of a defined Gq receptor (M3q), and stimulation of the endogenous Gq-coupled angiotensin receptor AT1 as parallel routes to manipulate PIP_2_. This design allows us to test directly whether depletion of the lipid itself alters Na_V_1.5 gating and to distinguish primary PIP_2_ effects from downstream DAG/PKC- and Ca² -dependent pathways.

We find that acute PIP_2_ loss, regardless of how it is achieved, destabilizes Na_V_1.5 gating, repositions both activation and steady-state inactivation, slows fast inactivation, and increases I_Na,L_. The effects are rapid, reversed by PIP_2_ enrichment, and independent of intracellular Ca² or PKC activity, indicating that membrane PIP_2_ is a direct and dominant determinant of Na_V_1.5 stability. Induced-fit docking and molecular dynamics simulations identify a PIP_2_-coupled pocket at the interface between the domain IV voltage sensor and pore that contains residues associated with proarrhythmic phenotypes, including R1644. Our *in silico* and electrophysiological studies reveal that the disease-reported variant R1644C (14, 15) weakens and redistributes predicted PIP_2_ contacts and shows a marked right shift in the apparent functional PIP_2_ sensitivity of late-current rescue. Together, these results provide a mechanistic framework linking lipid signaling, disease-associated variants, and altered cardiac excitability through PIP_2_-dependent stabilization of Na_V_1.5.

## Results

### Gq-PLC signaling shifts Na_V_1.5 gating through PIP_2_ depletion

Physiologically, PIP_2_ is rapidly hydrolyzed by PLC following activation of Gq-coupled GPCR signaling (16). To determine whether physiological Gq-PLC signaling modulates Na_V_1.5 through PIP_2_ depletion, we first activated endogenous angiotensin II type 1 (AT1) receptors in human iPSC-derived cardiomyocytes (iPS-CMs). Fig. 1A summarizes the AT1-Gq-PLC pathway and the pharmacological tools used to isolate PIP_2_-dependent signaling: candesartan, a selective AT1 receptor antagonist used to block Ang II receptor activation; the active PLC inhibitor U73122; the inactive analog U73433; the broad-spectrum kinase inhibitor staurosporine; and intracellular BAPTA to buffer Ca² .

**Figure 1.**
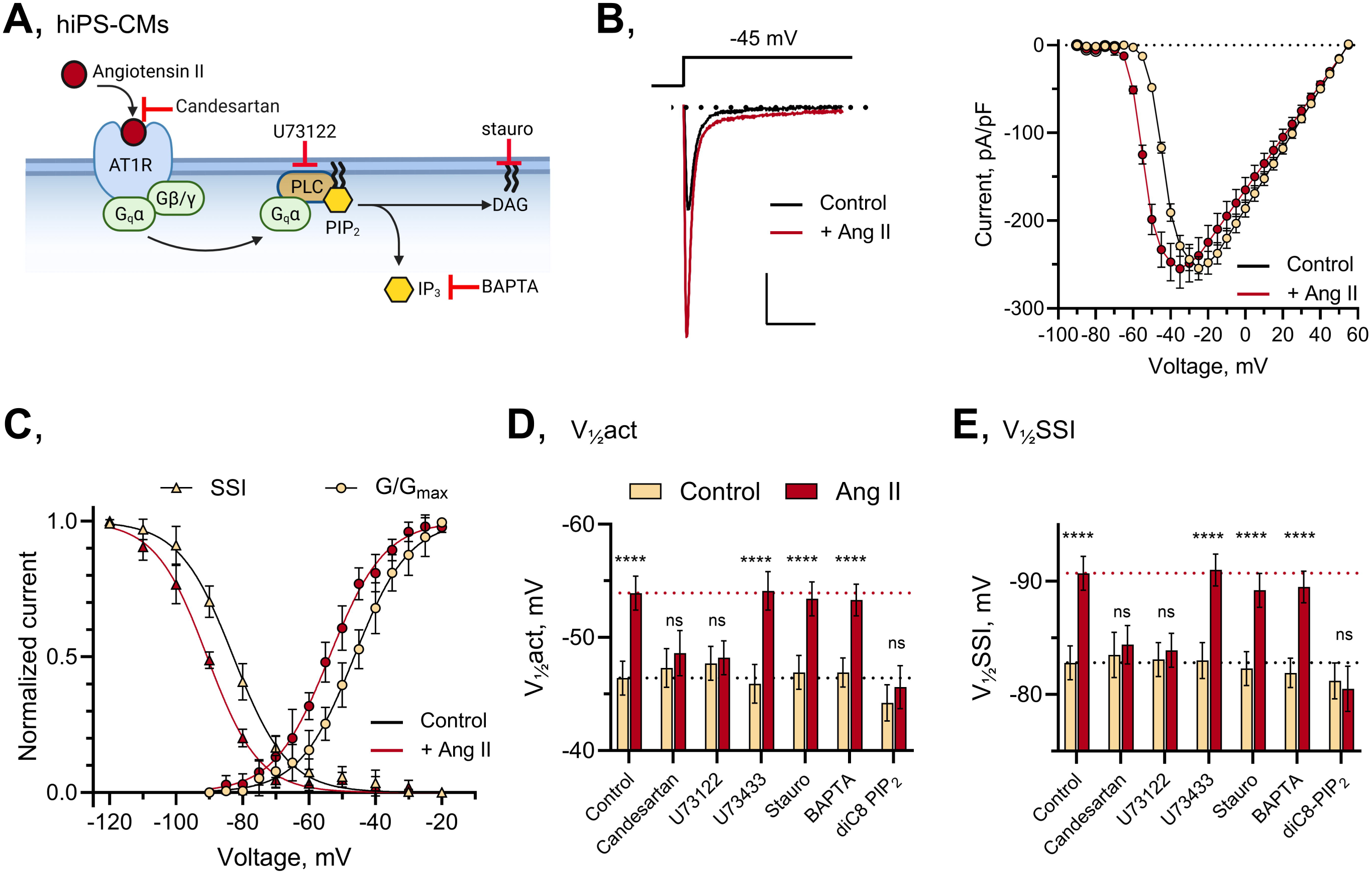
Gq-PLC signaling shifts Na_V_1.5 gating through PIP_2_ depletion. **(A)** Schematic of AT1 receptor activation and pharmacological strategy used to distinguish PLC-dependent PIP_2_ depletion from downstream PKC- or Ca^2+^-dependent signaling in human iPSC-derived cardiomyocytes. **(B)** Representative whole-cell I_Na_ traces (left), and mean current-voltage relationships (right) recorded before and after 100 nM angiotensin II (Ang II). **(C)**, Mean conductance-voltage relationship, and steady-state inactivation relationship showing Ang II-induced hyperpolarizing shifts in activation and availability. (**D, E**) Summary of V_½_act and V_½_SSI across pharmacological and lipid-loading conditions. The AT1 antagonist candesartan, U73122, and intracellular 50 µM diC8-PIP_2_ prevent the Ang II response, whereas U73433, staurosporine, and BAPTA do not. Values are means ± SD from 12–22 cells per condition, collected from at least 3 independent biological replicates. For direct comparisons in which currents were recorded from the same cell under two conditions, for example before and after Ang II perfusion, statistical significance was assessed using paired two-tailed t tests. Statistical symbols indicate comparisons of V½ values; ****p < 0.0001 versus the corresponding unstimulated control condition. See also Table S1.

Total internal reflection fluorescence microscopy (TIRFM), which restricts excitation to the plasma-membrane-adjacent evanescent field, confirmed that application of 100 nM angiotensin II (Ang II) decoupled the PIP_2_ biosensor iRFP-PH_PLCδ1_ from the plasma membrane of iPS-CMs with kinetics comparable to the optogenetic PIP_2_ depletion experiments described below (**Fig. S1**). Ang II-induced iRFP-PH_PLCδ1_ uncoupling was prevented by pretreatment with U73122 but not by U73433, supporting activation of a PLC-dependent Gq-signaling pathway in iPS-CMs.

Application of Ang II increased Na_V_1.5 current at subthreshold depolarizations and produced a leftward shift of approximately 10 mV in the current-voltage (I-V) relationship (**Fig. 1B**). For gating analysis, I-V data were converted to conductance, and normalized conductance-voltage (G-V) and steady-state inactivation (SSI) relationships were fitted with Boltzmann functions to obtain half-maximal voltages and slope factors. This analysis showed that Ang II caused parallel hyperpolarizing shifts in V½act (−7.5 mV; P < 0.001) and V½_SSI_ (−7.9 mV; P < 0.001), without significant changes in slope factors (**Fig. 1C, Table S1**). Thus, activation of endogenous AT1-Gq-PLC signaling repositions both activation and inactivation of native Na_V_1.5 in cardiomyocytes.

The Ang II-induced shifts in both V½act and V½_SSI_ were abolished by candesartan (300 nM) and by U73122 and were prevented when 50 µM diC8-PIP_2_ was included in the patch pipette. In contrast, U73433 did not prevent the Ang II-induced shifts, supporting a specific requirement for PLC activity rather than nonspecific inhibitor effects. Neither staurosporine (1 µM) nor intracellular BAPTA (10 mM) blocked the Ang II-induced shifts (**Fig. 1D, E, Table S1**). These data indicate that Ang II acts through AT1-Gq-PLC-dependent PIP_2_ hydrolysis to modulate Na_V_1.5 gating and argue against a dominant requirement for downstream PKC activation or changes in intracellular Ca² levels.

As an orthogonal test of Gq-PLC-dependent regulation, we expressed M3q, a Gq-coupled designer receptor exclusively activated by designer drugs (DREADD), in iPS-CMs and activated it with clozapine-N-oxide (CNO) (17). In the same TIRFM assay, application of CNO (100 nM) decoupled iRFP-PH_PLCδ1_ from the membrane interface of M3q-expressing iPS-CMs with kinetics like Ang II. CNO-induced iRFP-PH_PLCδ1_ uncoupling was prevented by pretreatment with U73122 but not by U73433 (**Fig. S1**). Using the same Boltzmann analysis, activation of M3q also produced gating shifts closely matching those observed with Ang II: V½act and V½_SSI_ each shifted by approximately 10 mV in the hyperpolarizing direction. This response was blocked by U73122 and by intracellular diC8-PIP_2_, but not by U73433, staurosporine, or BAPTA. As expected for this orthogonal receptor-ligand pair, candesartan had no effect on the M3q-mediated response to CNO (**Fig. S2**).

Together, these findings demonstrate that activation of Gq-PLC-coupled receptors in cardiomyocytes, whether endogenous AT1 receptors or an engineered Gq-DREADD, produces Na_V_1.5 gating changes that require PLC activity and are prevented by PIP_2_ loading. The pharmacological profile supports PLC-mediated PIP_2_ hydrolysis, rather than downstream PKC or Ca² signaling, as the dominant mechanism.

### Acute optogenetic PIP_2_ depletion confirms direct PIP_2_ control of Na_V_1.5 gating

To test whether PIP_2_ depletion itself is sufficient to reproduce the Gq-dependent Na_V_1.5 phenotype, we combined whole-cell patch-clamp recordings with optogenetically driven dephosphorylation of PIP_2_. This approach uses blue-light activation of Cryptochrome 2 (CRY2) fused to the tandem 4- and 5-phosphatase construct pseudojanin (CRY2-PJ). Co-expression of CRY2-PJ with CIBN-CAAX targets the enzyme to the inner leaflet of the plasma membrane upon photoactivation (10, 18, 19). Pseudojanin depletes PIP_2_ by sequential conversion to PI(4)P and PI (**Fig. 2A**) (20). We and others have validated this system using real-time PIP_2_ biosensors and canonical PIP_2_-sensitive channel readouts, including K_IR_, TMEM16A, and Na_V_1.4 currents (10, 21, 22).

**Figure 2.**
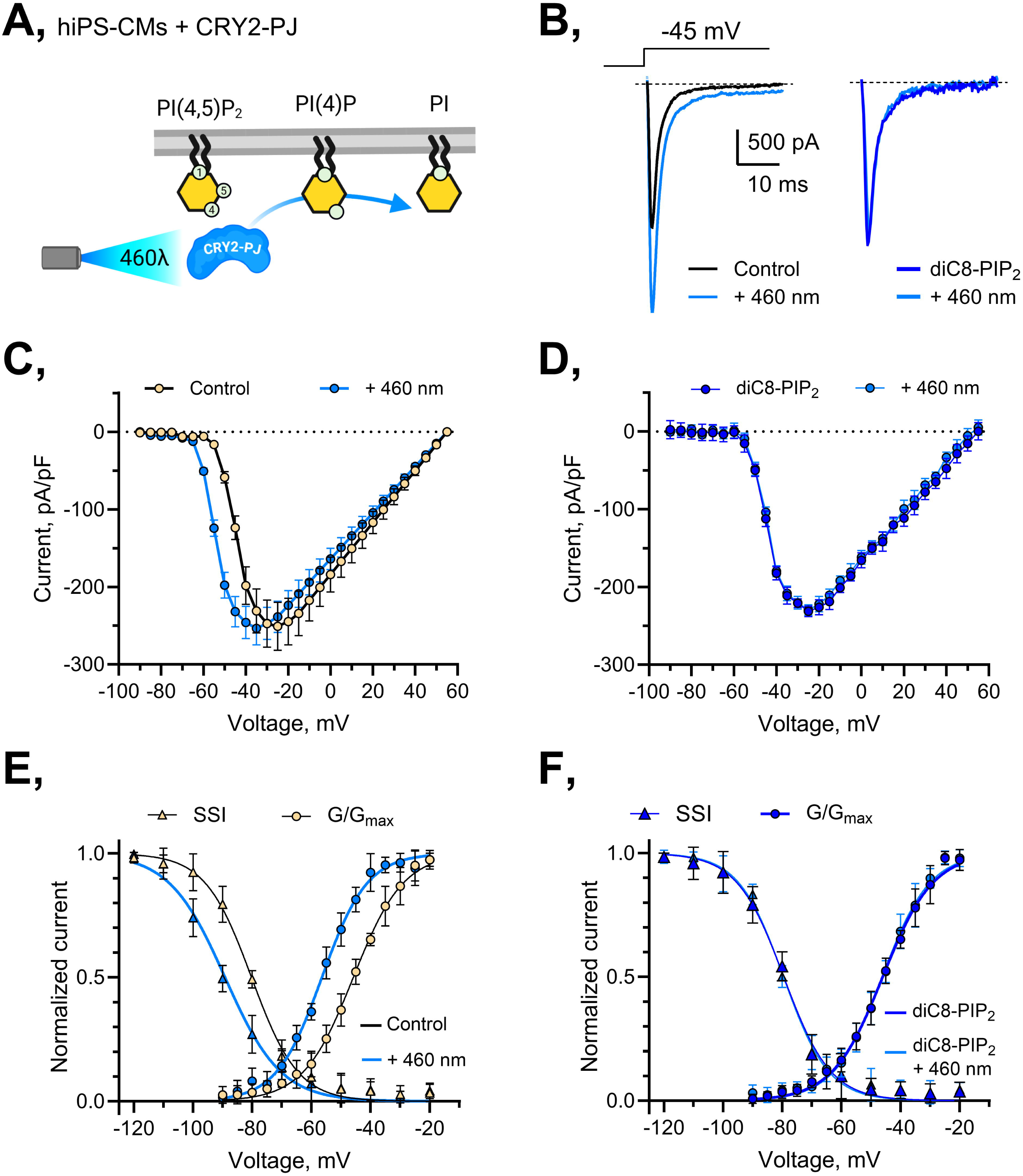
Acute optogenetic PIP_2_ depletion shifts Na_V_1.5 activation and inactivation in human iPSC-derived cardiomyocytes. **(A)** Schematic of CRY2-pseudojanin/CIBN-CAAX recruitment to the plasma membrane and enzymatic conversion of PIP_2_ to PI(4)P and PI after blue-light exposure. **(B)** Representative I_Na_ traces from iPS-CMs expressing CRY2-PJ/CIBN-CAAX before and after 460 nm (blue light) illumination, with or without 50 µM intracellular diC8-PIP_2_. (C, D) Current-voltage relationships showing that CRY2-PJ photoactivation enhances current at subthreshold voltages and shifts peak current to more negative potentials; intracellular diC8-PIP_2_ prevents this response. (**E, F**) Conductance-voltage and steady-state inactivation relationships showing coordinated hyperpolarizing shifts in V_½_act and V_½_SSI after PIP_2_ depletion. Values are means ± SD from 12–22 cells per condition, collected from at least 3 independent biological replicates. See also Table S1.

First, we monitored decoupling of the PIP_2_ biosensor iRFP-PH_PLCδ1_ from the plasma membrane of iPS-CMs following photoactivation of CRY2-PJ. TIRFM showed that iRFP-PH_PLCδ1_ rapidly decoupled from the membrane in response to blue-light illumination and was largely depleted from the membrane interface within approximately 200 s (**Fig. S1**). These findings are consistent with our prior observations and the work of others using iRFP-PH_PLCδ1_ in transfected tissue culture cells (10, 23–25).

Photoactivation of CRY2-PJ enhanced native whole-cell Na_V_1.5 current in iPS-CMs at submaximal depolarizations, especially near −40 mV, consistent with a hyperpolarizing shift in activation. Under control conditions, Na_V_1.5 currents activated near −55 mV and reached a peak current density of −250 ± 31 pA/pF at −25 mV. After CRY2-PJ photoactivation, the current-voltage relationship shifted leftward by approximately 10 mV, such that robust activation was observed at more negative voltages and peak current occurred near −35 mV without a substantive change in maximal current density (**Fig. 2B,C**). In cells recorded with 50 µM diC8-PIP_2_ in the patch pipette, blue light did not alter the I-V relationship or peak current density (**Fig. 2B,D**), supporting the conclusion that the effect of CRY2-PJ reflects PIP_2_ depletion rather than nonspecific photostimulation or optogenetic-component effects.

Using the same conductance conversion and Boltzmann fitting approach, we found that in control iPS-CMs, CRY2-PJ photoactivation caused an −11 ± 2 mV hyperpolarizing shift in the half-maximal activation voltage (V½act; P < 0.0001, paired t test) without a significant change in slope, and a parallel −10 ± 1 mV shift in V½_SSI_ (P < 0.0001, paired t test; **Fig. 2E,F**, **Table S1**). Thus, direct PIP_2_ depletion repositions both activation and inactivation toward more negative voltages. In cells dialyzed with 50 µM diC8-PIP_2_, neither V½act nor V½_SSI_ was altered by photoactivation of CRY2-PJ (**Fig. 2E,F, Table S1**).

Our prior Na_V_1.4 study showed that PIP_2_ depletion shifted V½act without substantively altering V½_SSI_ (10). To determine whether the coordinated activation and SSI shifts observed here reflect a cardiac-channel phenotype rather than a peculiarity of human iPS-CMs, we repeated CRY2-PJ/CIBN-CAAX photoactivation experiments in primary rat ventricular cardiomyocytes. CRY2-PJ produced quantitatively similar hyperpolarizing shifts in V½act and V½_SSI_, and these effects were abolished by 50 µM diC8-PIP_2_ in the pipette (**Fig. S3, Table S2**). Thus, Na_V_1.5 exhibits a distinct PIP_2_-dependent gating signature, with coordinated regulation of activation and inactivation that differs from our previous observations in Na_V_1.4.

### Convergent PIP_2_ depletion increases late Na_V_1.5 current

We next asked whether PIP_2_ depletion increases late Na_V_1.5 current. Because PIP_2_ depletion shifts both activation and SSI, steady-state voltage dependence alone does not uniquely predict the late-current phenotype. We therefore measured I_Na,L_ directly during long depolarizing steps to −30 mV delivered every 10 s and analyzed fast inactivation kinetics in the same recordings.

To relate late-current development to the kinetics of lipid depletion, we plotted I_Na,L_ over time during Ang II stimulation, M3q/CNO activation, and CRY2-PJ photoactivation and compared these traces with matched TIRFM measurements of iRFP-PH_PLCδ1_ membrane decoupling. I_Na,L_ developed over the same minutes-long interval in which iRFP-PH_PLCδ1_ was lost from the plasma membrane (**Fig. 3A-C; Fig. S1**). This temporal correspondence supports a direct relationship between membrane PIP_2_ depletion and late-current augmentation and provides a dynamic complement to the endpoint pharmacology shown below.

**Figure 3.**
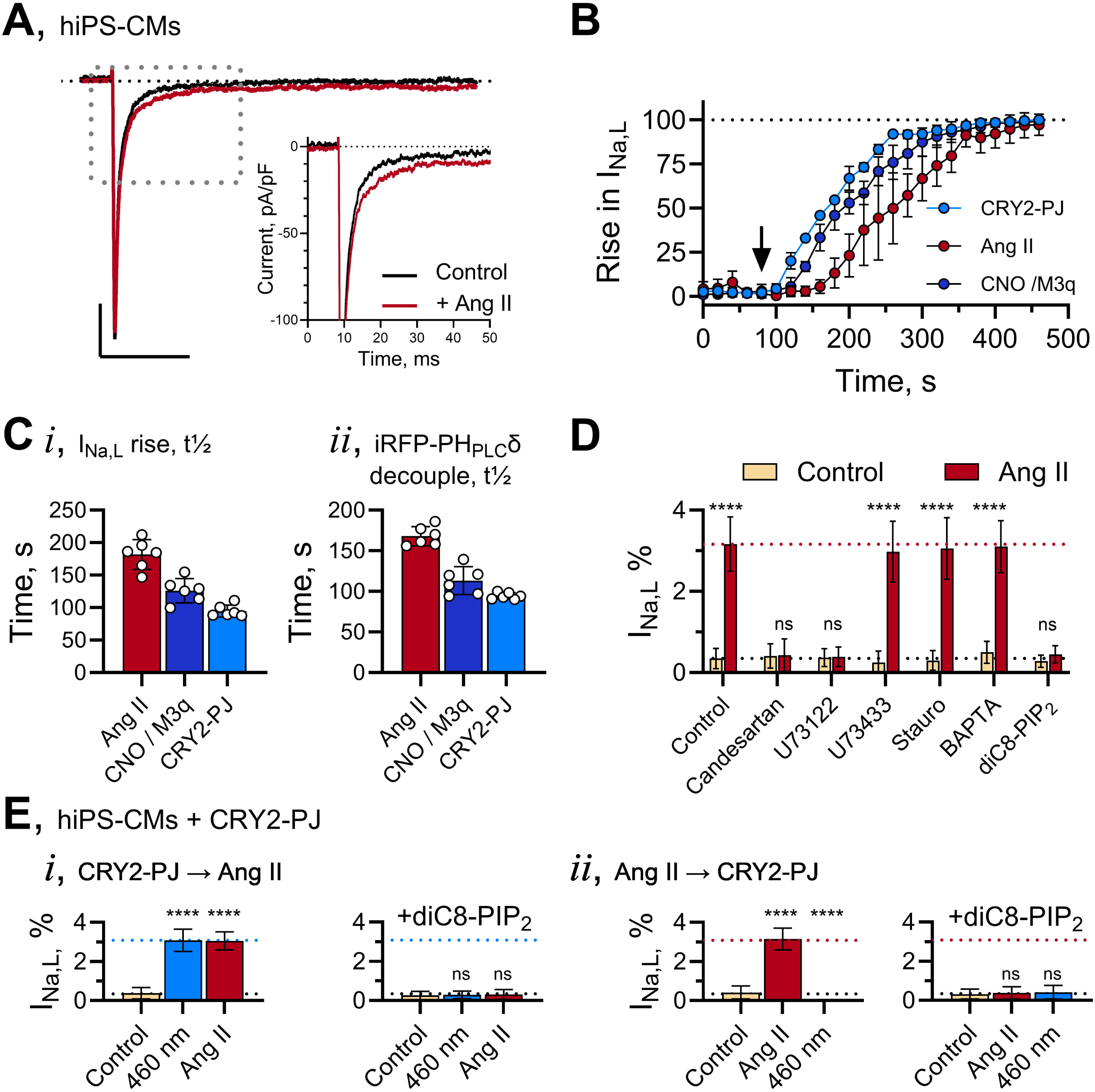
Convergent PIP_2_ depletion dynamically increases late Na_V_1.5 current. (A-C) Representative late-current recordings and time courses showing that Ang II, M3q/CNO signaling, and CRY2-PJ photoactivation each increase I_Na,L_ over a time course that parallels iRFP-PH_PLCδ1_ membrane decoupling. **(D)** Summary of Ang II-induced I_Na,L_ under control, antagonist, PLC-inhibitor, inactive-analog, kinase-inhibitor, Ca^2+^-buffering, and diC8-PIP_2_-loaded conditions. (**E, F**) Sequential activation experiments showing non-additivity between receptor-mediated and optogenetic PIP_2_ depletion. Values are means ± SD from 12–22 cells per condition, collected from at least 3 independent biological replicates. For direct comparisons in which currents were recorded from the same cell under two conditions, for example before and after Ang II perfusion, statistical significance was assessed using paired two-tailed t tests. Statistical symbols indicate comparisons of V½ values; ***p < 0.001 versus the corresponding unstimulated control condition. See also Table S3.

In control iPS-CMs, late current at −30 mV was small, comprising 0.35 ± 0.25% of the peak inward current. Application of 100 nM Ang II increased I_Na,L_ to 3.16 ± 0.67% of peak current (P < 0.001, paired t test; **Fig. 3A,D**). Ang II also slowed fast inactivation, increasing τ_fast_from 6.6 ± 0.9 ms under control conditions to 10.2 ± 1.1 ms (P < 0.01, paired t test). The Ang II-induced increase in I_Na,L_ and slowing of τ_fast_ were abolished by candesartan and U73122 and were prevented by 50 µM diC8-PIP_2_ in the patch pipette (**Fig. 3D**). In contrast, U73433, staurosporine, and BAPTA did not block the increase in late current or the slowing of fast inactivation. Thus, Ang II enhances I_Na,L_ through AT1-Gq-PLC signaling and PIP_2_ depletion, without a dominant requirement for downstream PKC activation or changes in intracellular Ca² (**Table S3**).

M3q activation produced a similar increase in I_Na,L_. In iPS-CMs expressing M3q, CNO increased late current to a level comparable to that produced by Ang II and produced a comparable slowing of τ_fast_; the pharmacological profile again matched a PIP_2_-dependent mechanism: the response was blocked by U73122 and intracellular diC8-PIP_2_, but not by U73433, staurosporine, or BAPTA. Candesartan did not prevent the M3q/CNO-induced increase in I_Na,L_ (**Fig. 3B, D; Table S3**). Representative M3q/CNO late-current traces and extended validation are shown in the supplement.

Finally, we directly compared optogenetic and receptor-mediated PIP_2_ depletion in the same cells. When CRY2-PJ was activated first by blue light, I_Na,L_ increased and τ_fast_ was prolonged to the same level observed with Ang II alone; subsequent application of Ang II produced no further effect **(Fig. 3E**). Conversely, when Ang II was applied first, late current reached its maximal level and subsequent CRY2-PJ photoactivation failed to augment it further. In both sequences, intracellular diC8-PIP_2_ prevented the increase in I_Na,L_. The ability of either manipulation to produce a full response, and the lack of additivity when the two are combined, supports convergence on a shared PIP_2_-sensitive mechanism.

Consistent with these cardiomyocyte findings, CRY2-PJ photoactivation also enhanced late Na_V_1.5 current in HEK293T cells expressing human Na_V_1.5, with a similar fold increase in I_Na,L_. As in iPS-CMs, the rise in late current was accompanied by a slowing of τ_fast_ in the same cells (control 7.0 ± 0.8 ms versus 12.4 ± 1.8 ms after exposure to blue light), and both effects were prevented by intracellular diC8-PIP_2_ (**Table S5**). Thus, in both human cardiomyocytes and a heterologous system, PIP_2_ depletion increases I_Na,L_ in parallel with a slowing of fast inactivation. Together, these results show that Gq-PLC-dependent PIP_2_ depletion, whether driven by endogenous AT1 receptors, an engineered Gq-DREADD, or optogenetic phosphatase recruitment, robustly increases late Na_V_1.5 current through a convergent membrane lipid mechanism.

### A DIV voltage sensor-pore PIP_2_ pocket provides a structural basis for Na_V_1.5 stabilization

To relate functional PIP_2_ dependence to specific structural interactions, we used induced-fit docking of PIP_2_ into a human Na_V_1.5 model based on PDB 7DTC (26). This analysis identified a PIP_2_-interaction pocket at the interface between the domain IV voltage sensor (DIV-VSD) and the adjacent pore domain (**Fig. 4A**). In the predicted wild-type pose, the PIP_2_ headgroup is coordinated by a cluster of basic side chains, with additional contacts contributed by nearby polar and hydrophobic residues. The resulting interaction-energy estimates support a stable PIP_2_ pose in this pocket but should be interpreted as relative structural scores rather than direct equilibrium binding free energies.

**Figure 4.**
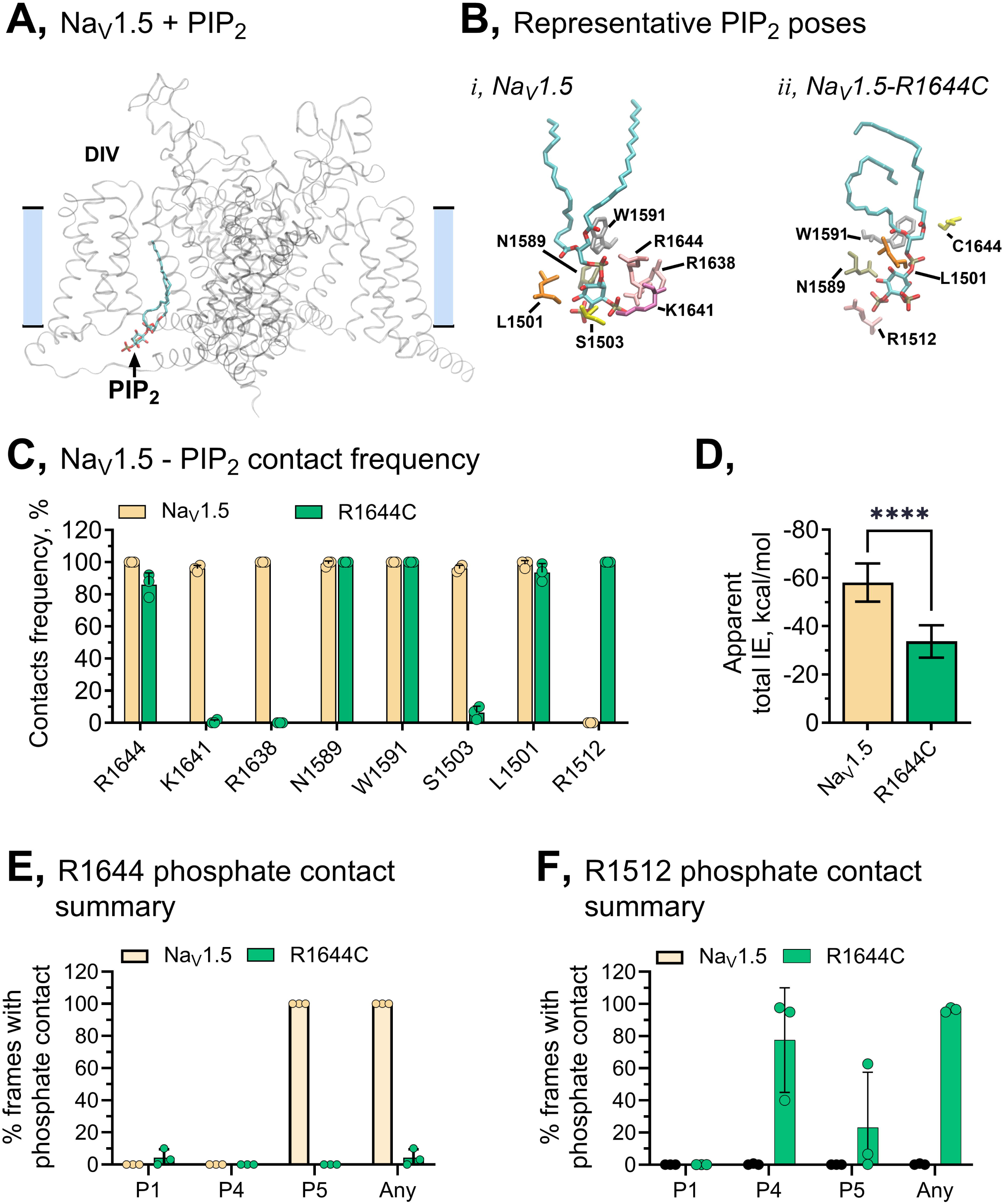
PIP_2_ occupies a DIV gating-interface pocket in Na_V_1.5 that is remodeled by R1644C. **(A)** Structural model showing the predicted PIP_2_-binding site at the interface between the domain IV voltage sensor and pore. **(B)** Representative PIP_2_ poses in wild-type Na_V_1.5 and R1644C after induced-fit docking and molecular dynamics simulation. **(C)** Contact-frequency analysis of residues contributing to the predicted pocket. **(D)** MM/GBSA estimates of apparent PIP_2_ interaction energy across independent replicate simulations. (**E, F**) Phosphate-specific contact analyses showing loss of R1644-phosphate contacts and emergence of R1512 contacts in the R1644C model. Data are mean ± SD where applicable.

Among the pocket residues, R1644 in the DIV S4-S5 linker, a residue altered in a reported inherited-arrhythmia variant, was the largest predicted contributor. Neighboring basic residues K1641 and R1638 also contributed substantially, so that the R1638-K1641-R1644 triad formed the core electrostatic cradle for the PIP_2_ headgroup. Additional contacts from N1589 and W1591 in DIV S5 helped define the pore-facing edge of the pocket (**Fig. 4B, C**). Thus, in the wild-type channel, PIP_2_ is predicted to occupy a compact interface between the DIV voltage sensor and pore, positioned to influence both voltage-sensor movement and coupling to inactivation.

Molecular dynamics simulations supported persistence of the predicted wild-type contact network over the analyzed trajectory windows (**Figs. S4; S5**). The minimum interaction distance between R1644 and PIP_2_ remained less than 5 Å in replicate simulations, with contacts occurring predominantly with the 5-phosphate on the inositol headgroup (**Fig. 4E; S5**). In contrast, the disease-reported R1644C variant retained PIP_2_ in the same general region but altered the pose and redistributed contacts. Loss of the positively charged guanidinium group abolished the strong R1644-centered interaction and reduced contacts with K1641 and R1638, whereas N1589 and W1591 continued to contribute to the binding scaffold (**Fig. 4B, C**). The weaker contact network was reflected in the binding energetics: molecular mechanics/generalized Born surface area (MM/GBSA) estimates of the apparent interaction energy were consistently less favorable for R1644C than for wild type across three independent replicas (**Fig. 4D**). Consistent with this structural shift, R1644-phosphate contacts were abolished in the R1644C system (**Fig. 4E**) (14, 15, 27, 28).

The R1644C model also promoted a new dominant interaction with R1512 in the DIV S2-S3 region. In simulations, the minimum distance between R1512 and the PIP_2_ headgroup decreased markedly relative to wild type, with contacts apparent with both the 4- and 5-phosphates (**Fig. 4C-F; Fig. S5**). Thus, R1644C is predicted not simply to weaken PIP_2_ coordination, but to redistribute the lipid from an R1644-centered electrostatic cradle toward a more dispersed network involving R1512. This structural prediction led us to test whether R1644C reduces functional PIP_2_-dependent stabilization of Na_V_1.5 late current.

### R1644C weakens functional PIP_2_-dependent stabilization of late current

We next tested the functional prediction that R1644C weakens PIP_2_-dependent stabilization. Because published functional characterization of this variant is limited, we interpreted R1644C primarily as a disease-reported perturbation of the predicted pocket and directly tested its PIP_2_-dependent phenotype under our recording conditions. In whole-cell recordings from HEK293T cells expressing wild-type Na_V_1.5, basal I_Na,L_ was small (0.35 ± 0.15% of peak current; peak current −323.5 ± 22 pA/pF; n = 45) and was not substantively changed by inclusion of diC8-PIP_2_ in the recording pipette (0.30 ± 0.10%; peak current −332.5 ± 20 pA/pF; **Fig. 5A, B, Table S5**). In contrast, Na_V_1.5-R1644C exhibited elevated basal I_Na,L_ (2.7 ± 0.2%), which was reduced to 1.1 ± 0.2% by diC8-PIP_2_. This rescue occurred without a substantive change in peak current density (−147.8 ± 37 pA/pF under control conditions versus −158 ± 41 pA/pF with diC8-PIP_2_; n = 15). Na_V_1.5-R1644A behaved similarly: I_Na,L_ was 3.2 ± 0.46% under control conditions and 1.35 ± 0.32% with 100 µM diC8-PIP_2_, without a significant change in peak current (**Fig. 5A,B; Tables S4 & S5**). Notably, basal τ_fast_ in R1644C was not slowed relative to wild-type channels under our recording conditions (**Table S5**). Thus, the elevated basal I_Na,L_ of R1644C is not explained simply by a slower macroscopic inactivation time constant, whereas acute PIP_2_ depletion slowed τ_fast_ and further increased I_Na,L_ in both wild-type and R1644C channels.

**Figure 5.**
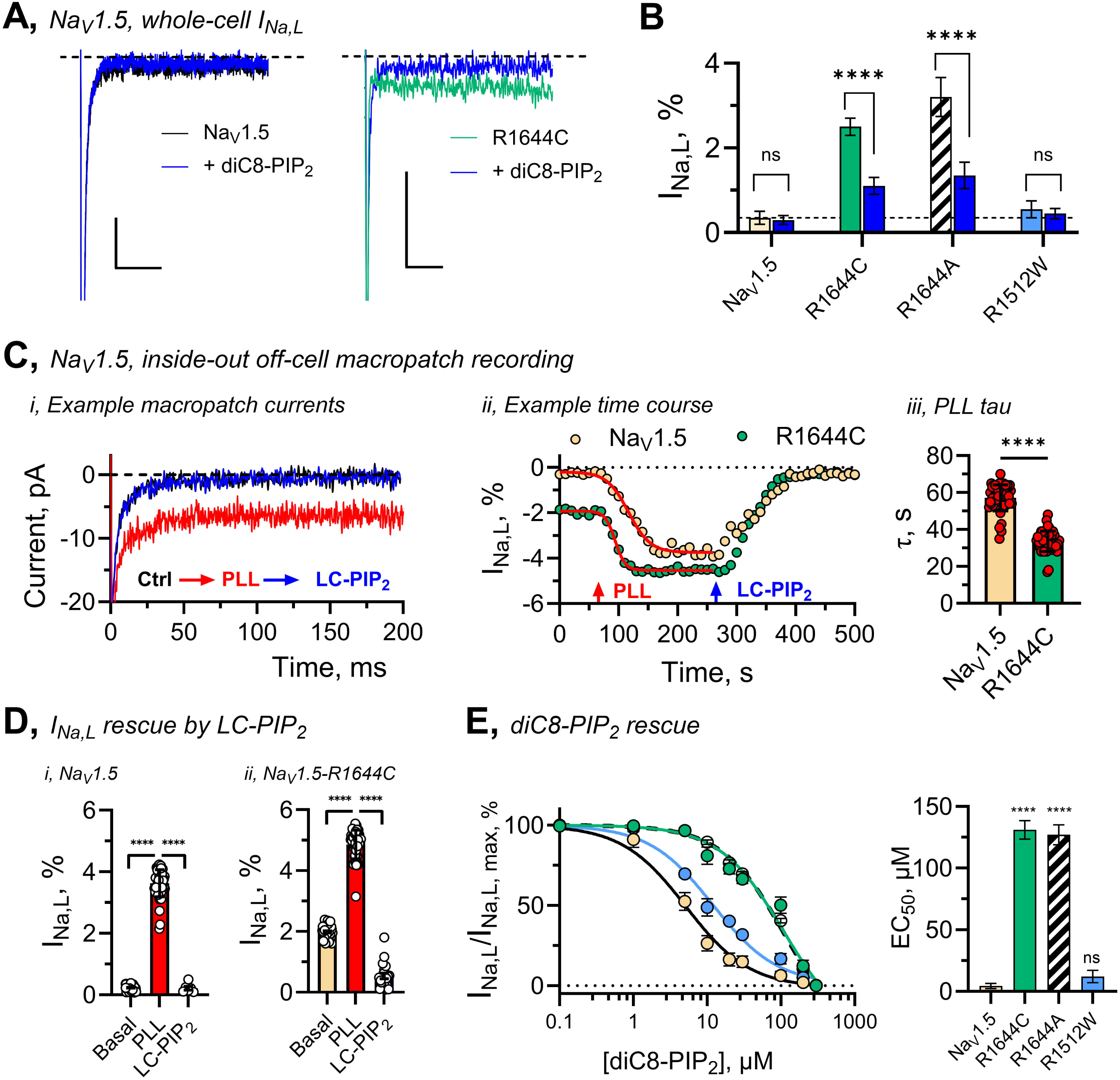
R1644C weakens PIP_2_-dependent suppression of late Na_V_1.5 current. **(A)** Representative whole-cell late-current traces from HEK293T cells expressing wild-type Na_V_1.5 or R1644C with or without intracellular 50 µM diC8-PIP_2_. **(B)** Summary of I_Na,L_ as a percentage of peak current for wild-type and mutant channels under control and diC8-PIP_2_-loaded conditions. **(C)** Inside-out macropatch experiment showing PLL-mediated functional depletion of membrane phosphoinositides and rescue by long-chain PIP_2_. **(D)** Summary of LC-PIP_2_ rescue of PLL- potentiated I_Na,L_ in wild-type Na_V_1.5 and R1644C patches. **(E)** Concentration-response analysis for diC8-PIP_2_ rescue of PLL-potentiated late current, showing a marked right shift in apparent functional PIP_2_ sensitivity for R1644C. Data are mean ± SD; n values and statistical comparisons are reported in Table S5.

Whole-cell rescue establishes that increasing intracellular PIP_2_ can suppress elevated late current in R1644C, but it does not provide a controlled measure of the lipid sensitivity of the channel at the membrane. To directly probe membrane-delimited PIP_2_ dependence, we used inside-out macropatches excised from HEK293T cells expressing either wild-type Na_V_1.5 or R1644C. Patches were held at 120 mV and pulsed to 30 mV every 10 s while late current was monitored during sequential depletion and replenishment of membrane PIP_2_ (**Fig. 5C**).

In wild-type Na_V_1.5 patches, basal I_Na,L_ was small, approximately 0.3% of peak current. Application of poly-L-lysine (PLL), which sequesters anionic phospholipids including PIP_2_ at the inner leaflet, produced a robust increase in late current to approximately 4% of peak. Subsequent application of 20 µM long-chain PIP_2_ (LC-PIP_2_) reversed this effect, returning I_Na,L_ toward basal levels (**Fig. 5C**). The time course of the PLL-induced increase in late current was well described by a single exponential with a time constant of approximately 60 s, consistent with gradual functional depletion of a membrane PIP_2_ pool.

R1644C patches behaved differently. Basal I_Na,L_ was already elevated before PLL application and increased further with PLL. Despite the higher basal and PLL-potentiated late current, LC-PIP_2_ still reversed the effect, restoring I_Na,L_ toward the pre-PLL level (**Fig. 5C,D**). The PLL-induced rise in late current was faster for R1644C than for wild-type channels, suggesting that the mutant channel is more sensitive to reductions in available PIP_2_.

To quantify relative PIP_2_ sensitivity, we next examined rescue of PLL-potentiated late current using defined concentrations of diC8-PIP_2_. Because aqueous diC8-PIP_2_ concentration does not directly report membrane mole fraction at the channel, these values are interpreted as apparent functional EC_50_ values rather than equilibrium binding constants. In wild-type patches, the diC8-PIP_2_ concentration-response relationship yielded an apparent EC_50_ of approximately 4 µM (**Fig. 5E**). In contrast, rescue of R1644C late current required much higher diC8-PIP_2_ concentrations, with an apparent EC_50_ of approximately 120 µM. Thus, R1644C shifts the apparent functional PIP_2_ sensitivity of Na_V_1.5 by approximately 30-fold, matching the structural prediction that the variant weakens and redistributes the PIP_2_-contact network.

These whole-cell and excised-patch data support the conclusion that R1644C weakens lipid-dependent stabilization rather than simply reducing channel expression or peak current. R1644C elevates basal I_Na,L_, responds more rapidly to PLL-mediated lipid sequestration, remains rescuable by LC-PIP_2_, and requires substantially higher diC8-PIP_2_ concentrations for functional suppression of late current. The convergence of these readouts argues that R1644C moves Na_V_1.5 toward a destabilized, PIP_2_-sensitive gating regime.

### R1512W perturbs channel availability without reproducing the R1644C late-current phenotype

Because the R1644C model predicted recruitment of R1512 as an alternative PIP_2_-contact residue, we tested whether perturbation of this node produces a related but distinct PIP_2_-sensitive phenotype. In whole-cell recordings from HEK293T cells, R1512W generated measurable peak currents but with lower current density than wild-type Na_V_1.5 under our recording conditions and exhibited a hyperpolarizing shift in steady-state availability with minimal effect on activation (**Tables S4 & S5**). The R1512W mutation is associated with Brugada and sudden unexplained nocturnal death syndrome in patients (29, 30). Intracellular diC8-PIP_2_ shifted the availability curve toward wild-type values, indicating that this gating defect remains lipid sensitive. Unlike R1644C, however, R1512W did not produce a prominent late-current phenotype under our conditions, and diC8-PIP_2_ did not significantly reduce I_Na,L_ (**Fig. 5A, B; Table S5**). Thus, R1512W should not be interpreted as a phenocopy of R1644C; rather, it supports the idea that neighboring nodes within the R1644C-redistributed PIP_2_-contact network can tune distinct aspects of Na_V_1.5 gating.

Together, these structural and functional results indicate that Na_V_1.5 relies on a PIP_2_-coupled network at the DIV-VSD-pore interface to constrain late openings and stabilize gating. R1644C weakens this network, increases basal and PIP_2_-depletion-evoked late current, and markedly reduces apparent functional PIP_2_ sensitivity. R1512W supports the broader conclusion that neighboring nodes within the same redistributed PIP_2_-contact network can bias Na_V_1.5 toward a different disease-relevant defect, primarily altered availability rather than a large late-current phenotype under our conditions.

## Discussion

This study identifies membrane PIP_2_ as a dynamic regulator of Na_V_1.5 gating and as a direct lipid intermediate linking Gq-PLC signaling to late sodium current in cardiomyocytes. Using endogenous AT1 receptor activation, engineered M3q signaling, optogenetic PIP_2_ depletion, intracellular and excised-patch lipid rescue, structural modeling, and disease-associated channel variants, we show that acute PIP_2_ depletion shifts Na_V_1.5 voltage-dependent gating, slows fast inactivation, and increases I_Na,L_. These effects are prevented or reversed by restoring PIP_2_, require PLC activity when triggered by Gq-coupled receptors, and do not require downstream PKC or Ca^2+^ signaling. Structural modeling places PIP_2_ in a pocket at the DIV voltage sensor-pore interface, where the disease-reported variant R1644C weakens and redistributes lipid contacts and shifts the apparent functional PIP_2_ sensitivity of late-current rescue by approximately 30-fold. Together, these data are most consistent with a model in which PIP_2_ stabilizes the inactivated machinery of Na_V_1.5 and thereby restrains proarrhythmic persistent sodium current.

### PIP_2_ acts within the modern allosteric model of fast inactivation

The interpretation of these findings is sharpened by the current structural view of sodium-channel fast inactivation. Classical models progressed from the Hodgkin-Huxley h-gate to ball-and-chain and hinged-lid mechanisms; more recent cryo-EM, fluorometry, gating-current, and mutagenesis studies support a more allosteric picture in which the IFM motif binds a receptor-like site that promotes closure of an intracellular inactivation gate rather than directly plugging the pore (31). In this framework, fast inactivation reflects coordinated motion among the DIV voltage sensor, the DIII-DIV linker, the S4-S5 linkers, and the intracellular ends of the S6 helices.

Our data place PIP_2_ within this framework rather than alongside it. PIP_2_ depletion does not behave like relief of a pore blocker; instead, the lipid tunes several coupled outputs of Na_V_1.5 gating at once: voltage-dependent activation and availability, the rate of fast inactivation, and the magnitude of I_Na,L_. This pattern is the signature expected if PIP_2_ stabilizes conformations that favor rapid and complete inactivation. Notably, recent multiscale simulations of Na_V_1.4 and Na_V_1.7 reach the same conclusion from an independent direction: PIP_2_ binds the inactivated channel and stabilizes the IFM (and an engineered IQM) motif within its receptor pocket, directly supporting a model in which the lipid reinforces the allosteric coupling that closes the inactivation gate (32). Because DIV voltage-sensor movement is closely linked to fast inactivation, the localization of the predicted PIP_2_ pocket at the DIV voltage sensor-pore interface is consistent with this role. We emphasize that we do not directly measure DIV sensor movement, IFM engagement, or inactivation-gate closure; the proposed link should therefore be read as a mechanistic framework to be tested rather than a structural observation.

### A conserved DIV voltage-sensor/S4-S5 PIP_2_ site shared across Na_V_ channels

Our induced-fit docking and molecular dynamics place PIP_2_ in a pocket at the interface between the DIV voltage sensor and the pore, coordinated by a triad of basic residues in the DIV S4-S5 linker (R1638, K1641, R1644) and supported by N1589 and W1591 in DIV S5. This assignment converges with an unbiased computational prediction made in the skeletal-muscle isoform. In coarse-grained and atomistic simulations of Na_V_1.4, PIP_2_ binds the inactivated channel at a DIV S4-S5 linker site (R1460, R1463, K1466, R1469) together with the DIII-DIV linker, and the same site is predicted to be conserved across Na_V_1.1-1.9 (22). The dominant coordinating residue in that model, R1469, is the structural analog of Na_V_1.5 R1644, the residue we identify as the largest predicted contributor to the wild-type pose and the one whose disease-associated substitution we characterize functionally. Independent sequence- and structure-based mapping had already flagged R1644C/H as the cardiac disease variant at this conserved position (33). Our excised-patch data, showing that R1644C markedly reduces apparent functional PIP_2_ sensitivity, therefore provide a direct experimental test, in the cardiac isoform, of a site predicted by unbiased simulation and conserved across the family.

That the same DIV S4-S5/voltage-sensor locus is implicated by induced-fit docking in Na_V_1.5, by unbiased sampling in Na_V_1.4 and Na_V_1.7, and by cross-channel sequence conservation argues against an idiosyncratic, isoform-specific interaction. The two computational approaches differ, however, in the auxiliary contacts they emphasize: the Na_V_1.4 simulations highlight DIII-DIV linker lysines and nearby DIV-region residues, whereas our Na_V_1.5 model implicates a pore-facing edge (N1589, W1591) and predicts redistribution toward R1512 in DIV S2-S3 when R1644 is lost. Notably, the Na_V_1.4 residue corresponding to Na_V_1.5 R1512 is R1337, which lies close to the predicted Na_V_1.4 PIP_2_-binding residue R1333 and was included in the RMSF analysis in that study. Thus, although R1512/R1337 should not be interpreted as an identical lipid-coordinating residue across the two models, the comparison supports the idea that the same DIV-adjacent structural neighborhood is dynamically coupled to PIP_2_ binding. Because our docking samples a single inactivated-state model (PDB 7DTC), whereas the Na_V_1.4 work samples binding and unbinding across long trajectories, we cannot yet determine whether the R1512 contact reflects a stable isoform-specific interaction, a state-dependent contact, or a redistribution favored by R1644C. Defining the pose in Na_V_1.5 by unbiased simulation and, ultimately, by direct structure determination remains an important goal.

### Isoform-specific lipid coupling between Na_V_1.4 and Na_V_1.5

PIP_2_ is well established as a cofactor for several potassium channels, and across voltage-gated channels more broadly it can tune voltage-sensor movement, pore opening, and inactivation, with a direction and consequence that are channel specific. PIP_2_ should therefore not be viewed as imposing a universal gating effect; local lipid contacts stabilize different conformational transitions according to each channel’s lipid-coupled gating interface.

Within the Na_V_ family, the available data now point to a shared site with isoform-specialized consequences. In Na_V_1.4, PIP_2_ acts as a negative regulator that stabilizes inactivation: its presence produces a depolarizing shift in activation, accelerates entry into fast inactivation, slows recovery from inactivation, and suppresses late current, while depletion has the opposite effect (10). Na_V_1.5 shares the central features, PIP_2_ depletion hyperpolarizes activation and increases late current, but differs in two respects: depletion produces coordinated hyperpolarizing shifts in both activation and steady-state inactivation, and it does so without a large change in peak current density. This points to isoform-specific lipid coupling on a conserved structural scaffold. We speculate that the more distributed contact network suggested by our Na_V_1.5 model, including the pore-facing residues and the R1512 region recruited in R1644C, could contribute to the broader, availability-coupled signature of the cardiac channel, although this remains a hypothesis. Isoform-specific cytoplasmic regulation of these two channels is itself well precedented: CaM-mediated Ca^2+^-dependent inactivation occurs in Na_V_1.4 but not Na_V_1.5 despite their homologous C-termini (33), making isoform-specialized PIP_2_ coupling mechanistically plausible rather than *ad hoc*.

The relationship between steady-state voltage shifts and I_Na,L_ is mechanistically linked but is not a simple one-to-one mapping. Because the activation and inactivation shifts are approximately parallel, changes in window current alone are unlikely to account for the several-fold increase in I_Na,L_. Across iPS-CMs and HEK293T cells, the rise in I_Na,L_ was accompanied by slowing of fast inactivation in the same recordings, consistent with destabilized inactivation contributing to persistent current beyond a steady-state window effect. Together with the temporal correspondence between PIP_2_ depletion and I_Na,L_ development, the reversible suppression of I_Na,L_ by diC8- or LC-PIP_2_, and the altered apparent PIP_2_ sensitivity of R1644C, these observations indicate that altered G-V/SSI relationships, slowed fast inactivation, and increased late current are complementary readouts of a common PIP_2_-dependent destabilization of Na_V_1.5 gating. Functionally this matters because even modest increases in I_Na,L_ can prolong the action potential, enhance Ca^2+^ entry, and promote afterdepolarizations.

### R1644C and the convergence of cytoplasmic inputs on late current

The disease-reported variant R1644C links pocket structure to function. Modeling predicts that R1644C abolishes the strong R1644-centered interaction, weakens contacts with K1641 and R1638, and shifts PIP_2_ toward an alternative network centered on R1512. Functionally, R1644C channels show elevated basal I_Na,L_, a faster response to PLL-mediated lipid sequestration, continued rescue by LC-PIP_2_, and a roughly 30-fold right shift in apparent diC8-PIP_2_ sensitivity. The convergence of these readouts argues that R1644C weakens lipid-dependent stabilization rather than simply reducing expression or peak current, moving Na_V_1.5 toward a destabilized, PIP_2_-sensitive gating regime.

This interpretation is reinforced by an independent biophysical signature of the variant. The conserved DIV S4-S5 PIP_2_ site is also where the C-terminal domain (CTD) is proposed to dock and sequester the DIII-DIV linker, dissociating to permit fast inactivation and rebinding during recovery; PIP_2_ is proposed to compete with the CTD at this site, so that reduced PIP_2_ coupling would be expected to accelerate recovery from inactivation (32, 34). Variants at this DIV S4–S5 locus have been associated with altered recovery from inactivation, and weakened PIP_2_ coordination at R1644C offers a testable framework for asking whether recovery from inactivation is also PIP_2_-sensitive. Weakened PIP_2_ coordination at R1644C offers a unifying explanation for that phenotype and predicts, as a direct and testable extension we did not measure here, that the accelerated recovery of R1644C is PIP_2_-sensitive.

More broadly, PIP_2_ appears to be one of several cytoplasmic inputs that converge on the DIV-coupled inactivation machinery to set late current. The CTD and calmodulin act through the same DIII-DIV linker/CTD axis: disruption of apo-CaM binding upregulates persistent Na_V_1.5 current, and CaM availability reverses it (33, 35). Our own work has shown that hypoxia drives proarrhythmic late current through SUMOylation of Na_V_1.5 (5). The present results add a lipid-based, Gq-coupled route to the same proarrhythmic endpoint, raising the possibility that distinct stressors, neurohumoral, metabolic, and genetic, converge on Na_V_1.5 late current through partly shared inactivation machinery.

Our data also refine an older view of how Gq signaling reaches cardiac and neuronal sodium channels. Modulation of Na_V_ current downstream of Gq-coupled receptors was historically attributed largely to PKC (11–13). Here, the same pharmacology that blocks the response to PLC inhibition leaves it intact under PKC inhibition and intracellular Ca^2+^ chelation, and intracellular PIP_2_ loading prevents it. These results reassign a major component of acute Gq modulation of Na_V_1.5 from a downstream kinase to direct depletion of the membrane lipid itself.

Finally, R1512 emerges as a neighboring node within the R1644C-remodeled DIV-adjacent network. In the wild-type model it contributes little to PIP_2_ coupling, but in R1644C simulations it becomes a dominant interacting residue. Experimentally, R1512W did not produce a prominent late-current phenotype under our conditions but did show a PIP_2_-sensitive shift in steady-state availability. R1512W should therefore not be read as a phenocopy of R1644C; rather, it supports the broader conclusion that neighboring residues in the same DIV-adjacent structural neighborhood can bias Na_V_1.5 toward distinct inactivation-related defects, with R1644C favoring late-current gain of function and reduced PIP_2_ sensitivity and R1512W primarily altering availability.

### Implications for receptor signaling, inherited variation, and excitability

The convergence of optogenetic and Gq-PLC manipulations on a shared PIP_2_-dependent mechanism has implications for how neurohumoral signals modulate cardiac excitability. Angiotensin II and other Gq-coupled agonists are elevated in heart failure, ischemia, and stress states associated with arrhythmia risk. Our results indicate that the action of PIP_2_ on the channel is membrane-delimited, since lipid depletion and rescue alter Na_V_1.5 late current in excised inside-out patches; the upstream receptor-to-lipid step, by contrast, requires intact Gq-PLC signaling. Together these define a route by which transient, receptor-driven PIP_2_ depletion can act locally at the channel to augment I_Na,L_, and the ability of PIP_2_ loading to prevent both AT1- and M3q-induced effects identifies local lipid availability as a key determinant of Na_V_1.5 behavior during Gq signaling.

The R1644C results extend this concept to inherited channel variation. By weakening PIP_2_-dependent stabilization and reducing apparent functional PIP_2_ sensitivity, R1644C places Na_V_1.5 closer to a destabilized regime in which basal late current is higher, lipid depletion produces a larger effect, and more PIP_2_ is required for rescue. Reduced PIP_2_ coupling may thus be a convergent mechanism through which genetic variation and acquired receptor signaling promote persistent sodium current. We frame the downstream consequences, effects on action-potential duration, conduction, and arrhythmia susceptibility, as predictions of this model to be tested in intact tissue and *in vivo*, rather than as outcomes established by the present recordings.

### Limitations and future directions

Several limitations warrant consideration. First, most experiments were performed in iPSC-derived cardiomyocytes and HEK293T cells, which do not fully recapitulate adult human myocytes; our neonatal rat ventricular myocyte data help bridge this gap, but intact tissue and *in vivo* models will be needed to define how PIP_2_-dependent Na_V_1.5 regulation affects action-potential parameters and arrhythmia. Second, our manipulations of PIP_2_ are relatively global and do not resolve microdomain-specific pools, such as those near intercalated discs or T-tubules, where Na_V_1.5 resides alongside distinct receptors and signaling partners.

Third, the structural analysis has defined limits. Our docking and molecular dynamics used a single, inactivated-state Na_V_1.5 cryo-EM model (PDB 7DTC) that lacks some peripheral and intrinsically disordered regions and does not capture all state-dependent conformations; only a subset of pocket residues has been probed experimentally, and some, such as K1641, may be essential for channel function and difficult to dissect by simple neutralization. Because we do not directly measure voltage-sensor movement, IFM engagement, or inactivation-gate closure, and because the related Na_V_1.4 and Na_V_1.7 simulations indicate that PIP_2_ interactions are state dependent (32), the proposed link between this pocket and the structural model of fast inactivation should be regarded as a framework to be tested. We cannot exclude additional PIP_2_-interaction surfaces, state-dependent lipid interactions, or accessory-protein-dependent contributions.

Finally, our lipid-rescue experiments use short- and long-chain PIP_2_ together with PLL-mediated sequestration. PLL sequesters anionic phospholipids broadly rather than PIP_2_ selectively, so the specificity of our conclusion rests on the PIP_2_ rescue rather than on the depletion step; and these approaches do not define the full lipid specificity of the pocket or the membrane mole fraction experienced by individual channels. Dissecting how different stimuli, including neurohumoral agonists beyond Ang II, metabolic stress, ischemia, and inherited variants, alter local PIP_2_ and related lipid pools, and whether R1644C alters recovery from inactivation in a PIP_2_-dependent manner, are important directions for future work.

## Conclusions

In summary, our data are consistent with a model in which Na_V_1.5 relies on a PIP_2_-coupled pocket at the DIV voltage sensor-pore interface to maintain stable inactivation and suppress late current. Acute PIP_2_ depletion, whether imposed optogenetically or through Gq-PLC-coupled receptors, shifts activation and availability, slows fast inactivation, and increases late sodium current in cardiomyocytes; restoring PIP_2_ reverses these effects. This site coincides with a PIP_2_-binding region independently predicted by unbiased simulation and conserved across the Na_V_ family, and the disease variant R1644C, at a residue analogous to a primary lipid-coordinating residue in Na_V_1.4, weakens lipid-dependent stabilization and markedly reduces apparent functional PIP_2_ sensitivity. Situated within current models of fast inactivation, these findings support the idea that PIP_2_ reinforces the DIV-coupled allosteric machinery that limits persistent Na^+^ conductance, and they position lipid signaling alongside CaM- and CTD-dependent regulation as convergent cytoplasmic inputs through which physiological signaling and inherited variation shape cardiac excitability and arrhythmia risk.

## Supplemental figure legends

**Figure S1. Total internal reflection fluorescence (TIRF) analysis of iRFP-PH_PLC_**_δ**1**_ **membrane decoupling in iPS-CMs.** Representative TIRF image series and summary time courses show loss of plasma-membrane iRFP-PH_PLCδ1_ fluorescence after Ang II stimulation, M3q/CNO activation, or CRY2-PJ photoactivation. U73122 prevents receptor-driven biosensor decoupling, whereas U73433 does not. Control experiments include unstimulated cells and optogenetic-component controls as indicated. Data are mean ± SD from the 6 – 8 cells per condition studies in three independent experiments.

**Figure S2. Gq-PLC signaling via the M3q DREADD shifts Na_V_1.5 gating through PIP_2_ depletion. (A)** Schematic of the M3q-DREADD/CNO strategy. **(B)** Representative iPS-CM I_Na_ traces before and after CNO. **(C)** Current-voltage, conductance-voltage, and steady-state inactivation relationships show CNO-induced hyperpolarizing shifts in activation and availability. **(D, E)** Summary analyses show that U73122 and intracellular diC8-PIP_2_ prevent the CNO response, whereas candesartan, U73433, staurosporine, and BAPTA do not. Data are mean ± SD; n values and statistical comparisons are reported in Tables S1 and S3.

**Figure S3. Acute PIP_2_ depletion shifts I_Na_ activation and inactivation in neonatal rat ventricular cardiomyocytes and HEK293T cells. (A-C)** Representative currents and summary gating analyses show that CRY2-PJ photoactivation produces coordinated hyperpolarizing shifts in V_½_act and V_½_SSI in neonatal rat ventricular myocytes, and that 50 µM intracellular diC8-PIP_2_ prevents these effects. Companion HEK293T experiments show a similar CRY2-PJ-dependent Na_V_1.5 gating phenotype in a heterologous system. Data are mean ± SD; n values and statistical comparisons are reported in Tables S2 and S4.

**Figure S4.** Root-mean-square deviation (RMSD) validation of Na_V_1.5-PIP_2_ molecular dynamics simulations. RMSD trajectories are shown for **(A)** three independent wild-type Na_V_1.5-PIP_2_ simulations and **(B)** three independent Na_V_1.5-R1644C-PIP_2_ simulations. These analyses support equilibration and trajectory stability over the production simulations used for contact and interaction-energy analyses.

**Figure S5.** Replicate molecular dynamics trajectories supporting PIP_2_-contact analysis. Replicate trajectories show minimum distances and atom contacts between PIP_2_ and key residues in the wild-type and R1644C systems, including R1644, R1512, and phosphate-specific contacts. Analyses support loss of R1644-centered contacts and redistribution toward R1512 in the R1644C model.

## Methods

### Molecular modelling and docking

The cryo-EM structure of human NaV1.5 (PDB ID: 7DTC) was prepared using the Protein Preparation Wizard module of Maestro (2021-1; Schrödinger, New York, NY) and subsequently used for docking. To systematically search potential PIP_2_-binding sites, the channel was covered with 56 grid boxes, each with centers spaced 20 Å apart, using an in-house script. Induced-fit docking (IFD) of the Glide program from Schrödinger was used for docking simulations (36, 37). diC1-PIP_2_ was initially placed at the center of each box for docking simulations using the standard precision (SP) module in Glide (38). Default parameters were used for IFD simulations. Residues within 3.5 Å of ligand poses were selected for side-chain optimization by Prime refinement. The SP score was used to rank ligand poses. For each potential binding site, the ligand pose with the lowest docking SP score was selected. Four predicted binding sites with the lowest docking SP scores were selected, and diC1-PIP_2_ was replaced with full-length PIP_2_ for MD simulations.

### System set-up and molecular dynamics simulations

Protonation states of the titratable residues in the Na_V_1.5 channel were calculated at pH = 7.4 via the use of the H++ server (http://biophysics.cs.vt.edu/) (39). The simulation system was generated using the CHARMM-GUI Membrane Builder webserver (http://www.charmmgui.org/?doc=input/membrane) (40). First, the protein was inserted in an explicit lipid bilayer of POPC, POPE, POPS and cholesterol with molecular ratio of 25:5:5:1 (41). Then, the complexes were put in a water box (145Å×145Å×132Å), followed by an addition of 150 mM NaCl to the system.

Molecular dynamics simulations were conducted using the PMEMD.CUDA program in AMBER 24. The tleap module was employed to neutralize the system by extra Na^+^ or Cl^−^ ions as needed. The FF19SB, LIPID21, and GAFF2 force fields were chosen for protein, mixed lipid membrane and PIP_2_, respectively. The parameters of PIP_2_ were generated using the general AMBER force field by the Antechamber module of AmberTools 25 and using the partial charge determined via restrained electrostatic potential charge-fitting scheme by ab initio quantum chemistry at the HF/6-31G* level. The systems were energetically minimized with 2,000 steps of steepest descent followed by 3,000 steps of conjugate gradient. Subsequently, Langevin dynamics was used to heat the systems from an initial temperature of 0 K to a final temperature of 303 K, with a collision frequency of 1 ps ¹. During heating, an initial constant force of 500 kcal·mol ¹·Å ² was applied to positionally restrain the receptor complexes, which was then gradually reduced to 10 kcal·mol ¹·Å ², allowing the lipid and water molecules to move freely. Subsequently, the systems underwent 5 ns of equilibrium molecular dynamics simulations. Finally, three independent 100-ns production molecular dynamics simulations were conducted for each system, with coordinates recorded every 100 ps for subsequent analysis. The simulations were conducted with periodic boundary conditions, initially in a constant temperature, constant-pressure ensemble (NPT), followed by a constant-temperature, constant volume ensemble (NVT). The pressure was regulated using the isotropic position scaling algorithm, with the pressure relaxation time set to 2.0 ps. The particle mesh Ewald (PME) method was used to calculate the long-range electrostatics with a 10 Å cut-off (42). An external voltage of 0.06 V/nm (approximately 200mV across the membrane) was added to the systems from the extracellular to the intracellular side, the treatment of the electric field has been detailed in previous work (43–47). A 4-fs time step was employed using the hydrogen mass repartition algorithm for systems to accelerate the molecular dynamics simulations (48).

### Molecular biology and reagents

Purified diC8-PIP_2_ and long-chain (LC) PIP_2_ were purchased from Avanti Biosciences. The LC-PIP_2_ (1-stearoyl-2-arachidonoyl-sn-glycero-3-phospho-(1’-myo-inositol-4’,5’-bisphosphate) is a synthetic PI(4,5)P_2_ with full length acyl chains. All PIP_2_ variants used in the patch-clamp studies were stored in sealed glass vials with low-binding caps and kept at −20 °C when not in use. In whole-cell recordings, diC8-PIP_2_ was included in the patch pipette at 50 µM for iPS-CM and rat ventricular myocyte experiments and at 50 or 100 µM for HEK293T experiments, as indicated. For excised inside-out patches, LC-PIP_2_ was applied to the cytoplasmic face of the membrane at a nominal concentration of 20 µM; because of the limited aqueous solubility of LC-PIP_2,_ this value is reported as an apparent concentration. All other reagents were purchased from Sigma or Tocris, unless otherwise noted.

Human Na_V_1.5 (NM_198056.1) was expressed from pcDNA1 together with human Na_V_1β (SCN1B channel, isoform b; GenBank: NM_001037), and iRFP, unless otherwise indicated, iRFP was used to identify transfected cells without photoactivation of optogenetic constructs. CIBN-CAAX was a gift from the De Camilli lab (Yale University, New Haven, CT). The open reading frame of pseudojanin (20) was designed with an N-terminal CRY2-tag in the pMAX(+) vector and was generated by Genscript (Piscataway, NY). DREADD-M3q was purchased from Addgene. Mutations were introduced by QuikChange mutagenesis according to the manufacturer’s protocols.

### Cell culture

Human embryonic kidney (HEK293T) cells were acquired from American Type Culture Collection (ATCC, Cat #CRL-3216) and were maintained in Dulbecco’s modified Eagle’s medium (ATCC) supplemented with 100 units/ml penicillin, 100 μg/ml streptomycin, and 10% (vol/vol) fetal bovine serum. Cells were routinely confirmed to be mycoplasma free via PCR and Hoechst stain. The cells were incubated in a 37°C, humidified incubator supplemented with 5% CO2. For experiments, cells were seeded on glass coverslips and transiently transfected with the indicated Na_V_1.5 constructs, Na_V_1β, iRFP, and, where indicated, CIBN-CAAX with CRY2-pseudojanin or the M3q receptor, in OptiMEM using polyethyleneimine (PEI) for 1-2 h at a ratio of 1 μg DNA to 4 μL PEI. iRFP was used as the transfection marker unless otherwise stated, allowing transfected cells to be identified without photoactivation of optogenetic constructs. Cells were studied 24-48 h after transfection.

Rat ventricular cardiomyocytes from neonatal Wistar rats were purchased from Lonza Biosciences. Cells were seeded at 300,000/cm2 on 1% fibronectin-coated coverslips or culture dishes (Thermo Fisher Scientific) and were maintained according to the manufacturer’s instructions in Bullet complete medium (Lonza) supplemented with 200 µM bromo-uridine to inhibit proliferation of cardiac fibroblasts.

Human induced pluripotent stem cell-derived cardiomyocytes (iPS-CMs) were purchased from Fujifilm Cellular Dynamics and cultured on #1.5 glass coverslips coated with 0.1% gelatin according to the manufacturer’s instructions and as described previously (5). Cells were maintained in iCell maintenance medium at 37°C in 5% CO / 95% air. iPS-CMs were studied between days 5 and 10 in culture, when isolated cells and syncytia exhibited rhythmic beating behavior. To enrich for NaV1.5-mediated current measurements, protocols and solutions were selected to minimize potassium and calcium conductances, and the recorded currents exhibited biophysical and pharmacological hallmarks described previously (5). For M3q-DREADD or CRY2 experiments, iPS-CMs were transfected with the indicated plasmids using Neuromag magnetic transfection (Oz Biosciences) 24-30 h before patch-clamp studies. Transfected cells were identified by iRFP fluorescence unless otherwise stated.

### Patch clamp recording

Currents were recorded with a Tecella Pico-2 amplifier (Tecella) controlled using WinWCP software (University of Strathclyde). Currents were low-pass Bessel filtered at 9 kHz and digitized at 50 kHz. Cells were studied in an external recording buffer comprising (in mM): 130 NaCl, 4 CsCl, 2 CaCl_2_, 1.2 MgCl_2_, 5 glucose, 10 HEPES, and 200 µM CdCl_2_, a concentration previously shown to block >98% of voltage-gated calcium channel current in primary cells (49). The pH was adjusted to 7.4 with NaOH. Patch pipettes were pulled from borosilicate glass (Clark Kent) using a vertical puller (Narishige) and had a resistance of 2.5-4 MΩ when filled with internal solution containing (in mM): 60 CsCl, 80 CsF, 1 MgCl_2_, 10 EGTA, and 5 Na_2_ATP, adjusted to pH 7.4 with CsOH. Pipettes were coated with Sylgard (Dow Corning) before use. Capacitance artifacts were subtracted online; series resistance was compensated to 70%, and cells with a series resistance of less than 10 MΩ were studied. Once whole-cell mode was established, cells were not studied for more than 600 s to maintain consistent membrane seals and voltage-clamp control of I_Na_. Current-voltage relationships were evoked from a holding potential of −100 mV by 100 ms test pulses between −100 and 60 mV, in 5 mV increments, unless otherwise stated. Steady-state inactivation was measured from a holding potential of −140 mV using a series of 100 ms conditioning prepulses from −120 to 20 mV, followed by a 50 ms test pulse to 0 mV to assess channel availability. A 10 s interpulse interval was used. Normalized peak current values are plotted against prepulse potential (mV). Activation relationships were fit with G/Gmax = 1/(1 + exp[(V1/2 - V)/k]), and SSI relationships were fit with I/Imax = 1/(1 + exp[(V - V1/2)/k]), where V1/2 is the voltage of half-maximal activation or inactivation and k is the slope factor. Whole-cell currents were normalized to cell capacitance. Mean ± SD capacitance values were 28 ± 5 pF for iPS-CMs and 12 ± 3 pF for HEK293T cells.

For excised (off-cell), inside-out patch experiments, pipettes were filled with the external recording buffer (in mM: 130 NaCl, 4 CsCl, 2 CaCl_2_, 1.2 MgCl_2_, 5 glucose, 10 HEPES, and 200 µM CdCl_2_, adjusted to pH 7.4 with NaOH), and the inside face of the patches was perfused with internal solution (in mM: 60 CsCl, 80 CsF, 1 MgCl_2_, 10 EGTA, and 5 Na_2_ATP, adjusted to pH 7.4 with CsOH). The polarity of the voltage-clamp protocols was inverted. To study concentration-dependent PIP_2_ rescue of channel activity, the current magnitude in the inside-out patch was measured, then the patch was perfused with poly-L-lysine (PLL; 100 µg/ml) to chelate phosphoinositides from the membrane. PIP_2_ variants (diC8- or LC-PIP_2_) were prepared in the internal recording buffer and applied to the inside face of the patch using a separate, vacuum-driven delivery system that terminates in a glass pipette placed in the recording chamber close to the patch. Although low-binding material tubing was used to deliver LC-PIP_2_, the final concentration of the reagent at the patch may be lower than the nominal concentration. Our readout for partition of the phospholipid into the membrane was a change in channel current. When using mutants with reduced or eliminated channel activity, parallel experiments with the wild-type channel were run to confirm that the reagent was active.

For simultaneous optogenetic patch-clamp studies, the blue-light system was photoactivated in epifluorescence mode using continuous excitation from a broad-spectrum LED (Excelitas) through a 488/20 nm filter (Chroma) via a 20x objective lens (Olympus). Light output at the sample was measured at 50 mW/cm2 by a photometer (ThorLabs). To avoid pre-activation of CRY2-fusion proteins, cells were incubated in the dark and handled in foil-wrapped dishes before experiments. Cells of interest were identified using iRFP fluorescence unless otherwise stated, avoiding blue-light exposure before the experiment. Further, a transilluminator that includes a bandpass filter was used to block blue light during brightfield visualization. To generate paired data, currents were recorded first in the dark and again following a 3-min pre-illumination with epifluorescent blue light that persisted through the remainder of the study. All experiments were performed at room temperature.

### Total internal reflection fluorescence (TIRF) microscopy

The PIP_2_ biosensor iRFP-PH_PLCδ1_ was imaged at the inner leaflet of live human iPS-derived cardiomyocytes or HEK293T cells by TIRF microscopy. Where indicated, cells co-expressed CRY2-pseudojanin and CIBN-CAAX, or the M3q-DREADD, together with iRFP-PH_PLCδ1_. Cells were seeded on #1.5 glass coverslips, transfected as described above, and studied after 24-48 hours in a bath solution comprising (in mM): 150 NaCl, 1 CaCl_2_, 1 MgCl_2_, 10 glucose, 10 HEPES, adjusted to pH 7.4 with NaOH. iRFP-PH_PLCδ1_ was excited by a 647-nm laser (OBIS, Coherent, Santa Clara, CA), and CRY2-fused phosphatases were photoactivated using a 445-nm laser. The excitation beams were conditioned for coherence using a custom-built Keplerian beam expander upstream of the laser clean-up filters. Each laser line was tuned to provide 10 mW of incident light on a micro mirror positioned below a high numerical-aperture apochromat objective (60x, 1.5 NA; Olympus, Waltham, MA) mounted on an RM21 microscope frame equipped with a piezo-driven nanopositioning stage (MadCity Labs, Madison, WI). The emission of iRFP-PH_PLCδ1_ was isolated from the excitation beam by an exit micro mirror and a ring-diaphragm positioned below the micro mirror assembly and imaged using a back-illuminated sCMOS camera (Teledyne Photometrics, Tucson, AZ) controlled by Micro-Manager freeware (UCSF). For receptor-stimulation experiments, 100 nM angiotensin II or 100 nM CNO was perfused into the bath, and image stacks were acquired at 5 s intervals with a 100 ms exposure to track membrane decoupling of the biosensor; where indicated, the PLC inhibitor U73122 or its inactive analog U73433 was applied before stimulation. All filters and mirrors were from Chroma (Bellows Falls, VT). Lenses, pinholes and diaphragms were from ThorLabs (Newton, NJ). TetraSpeck beads (Thermo, Waltham, MA) were routinely imaged to map the sCMOS chip and to calibrate the evanescent field depth to 100 nm (24). Images were acquired on at least three independent experimental days, with 3-4 coverslips analyzed per day; on each coverslip, 10-20 non-overlapping fields were imaged, and rectangular regions of interest (∼0.5-1 µm^2^) were drawn over the membrane footprint of individual cells. Laser power, exposure time, camera gain, and the 100 nm evanescent field depth were kept constant across all conditions within an experiment, and 30-60 cells were quantified per condition. The fluorescence intensity of cells was determined in ImageJ, with untransfected or unstimulated cells used to determine the background level of fluorescence.

### Statistics

Data were handled in WinWCP, Clampfit, Excel, and GraphPad Prism. Data are presented as mean ± SD unless otherwise indicated. Paired comparisons were analyzed using two-tailed paired t tests. For multi-condition pharmacology or mutant comparisons, statistical tests were selected according to experimental design, using repeated-measures or mixed-effects models with appropriate multiple-comparison correction where applicable. The threshold for significance was P < 0.05.

## Acknowledgements

This work was funded by an American Heart Association postdoctoral fellowship to K.D.G. (24POST1199293) and National Institutes of Health grant R01HL144615 to L.D.P. The authors thank Austin M. Baggetta, Anne K. Yauch, Takeharu Kawano, and Heikki Väänänen for technical support, and all members of the Plant laboratory for thoughtful discussion. CIBN-CAAX and iRFP-PH_PLCδ1_ were gifts from the De Camilli laboratory (Yale University, New Haven, CT). The computations were supported by the ITS (Information Technology Services) Research Computing at Northeastern University and the Argonne Leadership Computing Facility (ALCF) at Argonne National Laboratory

## Author Contributions

K.D.G, J.M.K., A.S.C., J.G.C., F.N., A.C., and L.D.P., Performed electrophysiological studies and data analysis. Z.M., X.M., Y.X., and M.C. performed computational studies and data analysis. K.D.G., performed TIRF microscopy and data analysis. K.D.G., A.C., and Y.X., designed and generated plasmid constructs. All authors contributed to writing the paper and preparing the figures. All authors commented on the manuscript.

## References

1. H. Abriel, Roles and regulation of the cardiac sodium channel Nav1.5: recent insights from experimental studies. Cardiovasc Res 76, 381–389 (2007).

2. L. D. Plant et al., A common cardiac sodium channel variant associated with sudden infant death in African Americans, SCN5A S1103Y. J Clin Invest 116, 430–435 (2006).

3. H. Abriel, J. S. Rougier, J. Jalife, Ion channel macromolecular complexes in cardiomyocytes: roles in sudden cardiac death. Circ Res 116, 1971–1988 (2015).

4. T. Kiyosue, M. Arita, Late sodium current and its contribution to action potential configuration in guinea pig ventricular myocytes. Circ Res 64, 389–397 (1989).

5. L. D. Plant, D. Xiong, J. Romero, H. Dai, S. A. N. Goldstein, Hypoxia Produces Pro-arrhythmic Late Sodium Current in Cardiac Myocytes by SUMOylation of NaV1.5 Channels. Cell Rep 30, 2225–2236 e2224 (2020).

6. L. S. Maier, S. Sossalla, The late Na current as a therapeutic target: where are we? J Mol Cell Cardiol 61, 44–50 (2013).

7. Y. Song, L. Belardinelli, Basal late sodium current is a significant contributor to the duration of action potential of guinea pig ventricular myocytes. Physiol Rep 5, e13295 (2017).

8. B.-C. Suh, B. Hille, PIP2 is a necessary cofactor for ion channel function: how and why? Annu. Rev. Biophys. 37, 175–195 (2008).

9. D. E. Logothetis et al., Phosphoinositide control of membrane protein function: a frontier led by studies on ion channels. Annu Rev Physiol 77, 81–104 (2015).

10. K. D. Gada et al., PI(4,5)P2 regulates the gating of NaV1.4 channels. J Gen Physiol 155 (2023).

11. J. Y. Ma, M. Li, W. A. Catterall, T. Scheuer, Modulation of brain Na+ channels by a G-protein-coupled pathway. Proc Natl Acad Sci U S A 91, 12351–12355 (1994).

12. A. R. Cantrell, J. Y. Ma, T. Scheuer, W. A. Catterall, Muscarinic modulation of sodium current by activation of protein kinase C in rat hippocampal neurons. Neuron 16, 1019–1026 (1996).

13. Y. Qu, J. Rogers, T. Tanada, T. Scheuer, W. A. Catterall, Modulation of cardiac Na+ channels expressed in a mammalian cell line and in ventricular myocytes by protein kinase C. Proc Natl Acad Sci U S A 91, 3289–3293 (1994).

14. C. Napolitano et al., Genetic testing in the long QT syndrome: development and validation of an efficient approach to genotyping in clinical practice. JAMA 294, 2975–2980 (2005).

15. A. Frustaci et al., Cardiac histological substrate in patients with clinical phenotype of Brugada syndrome. Circulation 112, 3680–3687 (2005).

16. B. Hille, E. Dickson, M. Kruse, B. Falkenburger, Dynamic metabolic control of an ion channel. Prog Mol Biol Transl Sci 123, 219–247 (2014).

17. B. N. Armbruster, X. Li, M. H. Pausch, S. Herlitze, B. L. Roth, Evolving the lock to fit the key to create a family of G protein-coupled receptors potently activated by an inert ligand. Proc Natl Acad Sci U S A 104, 5163–5168 (2007).

18. A. Chandrashekar et al., SUMOylation and an ATS1 variant converge to disrupt PIP2-dependent gating of Kir2.1. J Gen Physiol 157 (2025).

19. K. D. Gada et al., Optogenetic dephosphorylation of phosphatidylinositol 4,5 bisphosphate in Xenopus laevis oocytes. STAR Protoc 4, 102003 (2023).

20. G. R. Hammond et al., PI4P and PI(4,5)P2 are essential but independent lipid determinants of membrane identity. Science 337, 727–730 (2012).

21. O. Idevall-Hagren, E. J. Dickson, B. Hille, D. K. Toomre, P. De Camilli, Optogenetic control of phosphoinositide metabolism. Proceedings of the National Academy of Sciences of the United States of America 109, E2316–E2323 (2012).

22. J. Xu et al., Allosteric coupling between PIP(2) and Ca(2+) binding sites gates TMEM16A channels. Proc Natl Acad Sci U S A 123, e2506040123 (2026).

23. O. Idevall-Hagren, P. Decamilli, Manipulation of plasma membrane phosphoinositides using photoinduced protein-protein interactions. Methods Mol Biol 1148, 109–128 (2014).

24. K. D. Gada, J. M. Kamuene, T. Kawano, L. D. Plant, Imaging Membrane Proteins Using Total Internal Reflection Fluorescence Microscopy (TIRFM) in Mammalian Cells. Bio Protoc 13, e4614 (2023).

25. K. D. Gada, T. Kawano, L. D. Plant, D. E. Logothetis, An optogenetic tool to recruit individual PKC isozymes to the cell surface and promote specific phosphorylation of membrane proteins. J Biol Chem 298, 101893 (2022).

26. Z. Li et al., Structure of human Nav1.5 reveals the fast inactivation-related segments as a mutational hotspot for the long QT syndrome. Proc Natl Acad Sci U S A 118 (2021).

27. B. M. Kroncke, A. M. Glazer, D. K. Smith, J. D. Blume, D. M. Roden, SCN5A (Na(V)1.5) Variant Functional Perturbation and Clinical Presentation: Variants of a Certain Significance. Circ Genom Precis Med 11, e002095 (2018).

28. A. Frustaci, M. A. Russo, C. Chimenti, Structural myocardial abnormalities in asymptomatic family members with Brugada syndrome and SCN5A gene mutation. Eur Heart J 30, 1763 (2009).

29. I. Deschenes et al., Electrophysiological characterization of SCN5A mutations causing long QT (E1784K) and Brugada (R1512W and R1432G) syndromes. Cardiovasc Res 46, 55–65 (2000).

30. L. Zhang et al., Does Sudden Unexplained Nocturnal Death Syndrome Remain the Autopsy-Negative Disorder: A Gross, Microscopic, and Molecular Autopsy Investigation in Southern China. Mayo Clin Proc 91, 1503–1514 (2016).

31. Y. Liu et al., Evolution of our understanding of the sodium channel fast inactivation: From Hodgkin and Huxley to the structural era. J Gen Physiol 158 (2026).

32. Y. Lin, E. Tao, J. P. Champion, B. Corry, A binding site for phosphoinositides described by multiscale simulations explains their modulation of voltage-gated sodium channels. Elife 12 (2024).

33. S. Nathan et al., Structural basis of cytoplasmic NaV1.5 and NaV1.4 regulation. J Gen Physiol 153 (2021).

34. T. Clairfeuille et al., Structural basis of alpha-scorpion toxin action on Na(v) channels. Science 363 (2019).

35. H. Yan, C. Wang, S. O. Marx, G. S. Pitt, Calmodulin limits pathogenic Na+ channel persistent current. J Gen Physiol 149, 277–293 (2017).

36. W. Sherman, H. S. Beard, R. Farid, Use of an induced fit receptor structure in virtual screening. Chem Biol Drug Des 67, 83–84 (2006).

37. W. Sherman, T. Day, M. P. Jacobson, R. A. Friesner, R. Farid, Novel procedure for modeling ligand/receptor induced fit effects. J Med Chem 49, 534–553 (2006).

38. R. A. Friesner et al., Glide: a new approach for rapid, accurate docking and scoring. 1. Method and assessment of docking accuracy. J Med Chem 47, 1739–1749 (2004).

39. J. C. Gordon et al., H++: a server for estimating pKas and adding missing hydrogens to macromolecules. Nucleic Acids Res 33, W368–371 (2005).

40. S. Jo, T. Kim, V. G. Iyer, W. Im, CHARMM-GUI: a web-based graphical user interface for CHARMM. J Comput Chem 29, 1859–1865 (2008).

41. E. Leal-Pinto et al., Gating of a G protein-sensitive mammalian Kir3.1 prokaryotic Kir channel chimera in planar lipid bilayers. J Biol Chem 285, 39790–39800 (2010).

42. T. A. Darden, L. G. Pedersen, Molecular modeling: an experimental tool. Environ Health Perspect 101, 410–412 (1993).

43. X. Y. Meng, S. Liu, M. Cui, R. Zhou, D. E. Logothetis, The Molecular Mechanism of Opening the Helix Bundle Crossing (HBC) Gate of a Kir Channel. Sci Rep 6, 29399 (2016).

44. M. Cui et al., A novel small-molecule selective activator of homomeric GIRK4 channels. J Biol Chem 298, 102009 (2022).

45. D. Li, T. Jin, D. Gazgalis, M. Cui, D. E. Logothetis, On the mechanism of GIRK2 channel gating by phosphatidylinositol bisphosphate, sodium, and the Gbetagamma dimer. J Biol Chem 294, 18934–18948 (2019).

46. B. Roux, The membrane potential and its representation by a constant electric field in computer simulations. Biophys J 95, 4205–4216 (2008).

47. P. Bjelkmar, P. S. Niemela, I. Vattulainen, E. Lindahl, Conformational changes and slow dynamics through microsecond polarized atomistic molecular simulation of an integral Kv1.2 ion channel. PLoS Comput Biol 5, e1000289 (2009).

48. C. W. Hopkins, S. Le Grand, R. C. Walker, A. E. Roitberg, Long-Time-Step Molecular Dynamics through Hydrogen Mass Repartitioning. J Chem Theory Comput 11, 1864–1874 (2015).

49. H. A. Pearson, K. G. Sutton, R. H. Scott, A. C. Dolphin, Ca2+ currents in cerebellar granule neurones: role of internal Mg2+ in altering characteristics and antagonist effects. Neuropharmacology 32, 1171–1183 (1993).

