## Supplemental materials for "PIP_2_ stabilizes Na_V_1.5 gating and links receptor signaling to cardiac late sodium current"

**Tables**

**Table S1.** Voltage-dependent activation and steady-state inactivation parameters for $I_{Na}$ in human iPSC-derived cardiomyocytes.

**Table S2.** PIP$_2$-dependent regulation of Na$_V$ current gating and late sodium current in neonatal rat ventricular myocytes.

**Table S3.** Peak and late $I_{Na}$ parameters in human iPS-derived cardiomyocytes.

**Table S4.** Voltage dependence of activation and steady-state fast inactivation of WT and mutant Na$_V$1.5 channels expressed in HEK293T cells.

**Table S5.** Late sodium current and fast-inactivation kinetics of WT and mutant Na$_V$1.5 channels expressed in HEK293T cells.

**Supplemental figures:**

**Figure S1.** Total internal reflection fluorescence (TIRF) analysis of iRFP-PH$_{PLC\delta1}$ membrane decoupling in iPS-CMs.

**Figure S2.** Gq-PLC signaling via the M3q DREADD shifts Na$_V$1.5 gating through PIP$_2$ depletion.

**Figure S3.** Acute PIP$_2$ depletion shifts $I_{Na}$ activation and inactivation in neonatal rat ventricular cardiomyocytes and HEK293T cells.

**Figure S4.** Root-mean-square deviation (RMSD) validation of Na$_V$1.5-PIP$_2$ molecular dynamics simulations.

**Figure S5.** Replicate molecular dynamics trajectories supporting PIP$_2$-contact analysis.

**Table S1**

| Experiment | V½$_{act}$, mV | K$_{act}$, mV | V½$_{SSI}$, mV | K$_{SSI}$, mV |
|---|---|---|---|---|
| **Untransfected iPS-CMs** | | | | |
| Control | -46.4 ± 1.5 | 7.8 ± 0.6 | -82.8 ± 1.5 | 6.8 ± 0.6 |
| Control + candesartan | -47.3 ± 1.7 | 8.0 ± 0.5 | -83.5 ± 2.0 | 7.0 ± 0.5 |
| Control + U73122 | -47.7 ± 1.5 | 7.7 ± 0.6 | -83.1 ± 1.6 | 6.7 ± 0.8 |
| Control + U73433 | -45.9 ± 1.6 | 7.9 ± 0.4 | -83.0 ± 1.5 | 6.8 ± 0.6 |
| Control + Staurosporine | -46.9 ± 1.4 | 7.8 ± 0.5 | -82.3 ± 1.5 | 7.2 ± 0.7 |
| Control + BAPTA | -46.9 ± 1.3 | 7.8 ± 0.6 | -81.9 ± 1.3 | 6.5 ± 0.5 |
| Control + diC8-$PIP_2$ | -44.2 ± 1.6 | 7.9 ± 0.6 | -81.2 ± 1.6 | 6.4 ± 0.5 |
| Ang II | -55.9 ± 1.6$^{***}$ | 8.1 ± 0.5 | -91.7 ± 1.5$^{***}$ | 7.8 ± 1.1 |
| Ang II + candesartan | -48.6 ± 2.0 | 7.8 ± 0.8 | -84.4 ± 1.7 | 7.2 ± 1.2 |
| Ang II + U73122 | -48.2 ± 1.5 | 7.9 ± 0.7 | -83.9 ± 1.5 | 6.8 ± 0.8 |
| Ang II + U73433 | -54.1 ± 1.7$^{***}$ | 7.6 ± 0.9 | -91.0 ± 1.4$^{***}$ | 7.1 ± 0.6 |
| Ang II + Staurosporine | -53.4 ± 1.5$^{***}$ | 8.0 ± 1.0 | -89.2 ± 1.5$^{***}$ | 7.6 ± 0.6 |
| Ang II + BAPTA | -53.3 ± 1.4$^{***}$ | 7.9 ± 0.6 | -89.5 ± 1.5$^{***}$ | 7.3 ± 0.6 |
| Ang II + diC8-$PIP_2$ | -45.6 ± 1.9 | 7.8 ± 0.8 | -80.5 ± 2.0 | 6.8 ± 0.6 |
| **M3q-DREADD iPS-CMs** | | | | |
| Control | -46.5 ± 2.0 | 8.0 ± 1.2 | -81.4 ± 2.1 | 8.1 ± 0.9 |
| Control + candesartan | -47.1 ± 1.8 | 7.9 ± 0.7 | -81.2 ± 2.0 | 8.3 ± 1.1 |
| Control + U73122 | -46.9 ± 2.0 | 8.1 ± 0.8 | -80.8 ± 1.8 | 8.1 ± 0.8 |
| Control + U73433 | -47.0 ± 1.8 | 8.0 ± 1.1 | -82.0 ± 2.0 | 8.0 ± 0.8 |
| Control + Staurosporine | -46.5 ± 2.1 | 8.2 ± 1.0 | -80.9 ± 2.1 | 7.8 ± 0.9 |
| Control + BAPTA | -46.6 ± 2.0 | 7.8 ± 0.7 | -79.6 ± 2.0 | 8.1 ± 0.9 |
| Control + diC8-$PIP_2$ | -44.5 ± 1.8 | 8.1 ± 1.3 | -79.2 ± 2.2 | 7.9 ± 1.3 |
| CNO | -56.3 ± 2.2$^{***}$ | 8.4 ± 0.9 | -91.7 ± 2.3$^{***}$ | 8.2 ± 1.1 |
| CNO + candesartan | -56.1 ± 2.0$^{***}$ | 8.2 ± 1.2 | -89.9 ± 2.1$^{***}$ | 8.3 ± 1.0 |
| CNO + U73122 | -47.3 ± 2.1 | 8.1 ± 1.1 | -81.4 ± 2.1 | 8.2 ± 0.8 |
| CNO + U73433 | -56.8 ± 2.4$^{***}$ | 8.1 ± 1.2 | -92.0 ± 2.1$^{***}$ | 7.9 ± 0.9 |
| CNO + Staurosporine | -55.8 ± 2.1$^{***}$ | 8.0 ± 1.3 | -89.5 ± 2.0$^{***}$ | 7.8 ± 0.7 |

| | | | | |
|---|---|---|---|---|
| CNO + BAPTA | -56.0 ± 2.3$^{***}$ | 7.9 ± 0.9 | -89.1 ± 1.7$^{***}$ | 8.1 ± 0.8 |
| CNO + diC8-$PIP_2$ | -45.1 ± 2.1 | 8.3 ± 1.1 | -80.0 ± 2.6 | 8.0 ± 1.1 |
| **CRY2-PJ iPS-CMs** | | | | |
| Control, dark | -45.8 ± 1.9 | 8.2 ± 0.8 | -80.3 ± 2.2 | 7.8 ± 0.6 |
| Control + 460 nm | -56.5 ± 2.1$^{***}$ | 8.4 ± 0.9 | -90.0 ± 2.0$^{***}$ | 8.1 ± 0.9 |
| diC8-$PIP_2$, dark | -46.0 ± 2.1 | 8.5 ± 0.7 | -79.5 ± 2.0 | 8.0 ± 1.0 |
| diC8-$PIP_2$ + 460 nm | -46.3 ± 2.2 | 8.4 ± 0.5 | -79.7 ± 2.1 | 8.2 ± 1.1 |

**Table S1: Voltage-dependent activation and steady-state inactivation parameters for $I_{Na}$ in human iPSC-derived cardiomyocytes.** Whole-cell sodium currents were recorded from human iPS-derived cardiomyocytes under the indicated experimental conditions. Where indicated, cells were transfected to express the CNO-activated M3q-DREADD or light-activated CRY2-pseudojanin together with CIBN-CAAX. Voltage-clamp protocols are described in the Methods. Activation parameters were obtained from conductance-voltage relationships, where conductance was calculated from the current-voltage relationship and normalized to maximal conductance. Steady-state inactivation (SSI) parameters were obtained from normalized current availability relationships. Activation relationships were fit with a Boltzmann function, $G/Gmax = 1/(1 + \exp[(V_{½} - V)/k])$, and SSI relationships were fit with $I/Imax = 1/(1 + \exp[(V - V_{½})/k])$, where $V_{½}$ is the voltage of half-maximal activation or inactivation and k is the slope factor. Values are means ± SD from 12–22 cells per condition, collected from at least 3 independent biological replicates. The mean whole-cell capacitance value was 28 ± 5 pF. For direct comparisons in which currents were recorded from the same cell under two conditions, for example before and after Ang II perfusion, statistical significance was assessed using paired two-tailed t tests. For comparisons among multiple independent treatment groups, statistical tests are indicated in the table footnotes. Statistical symbols indicate comparisons of $V_{½}$ values; ***$p < 0.001$ versus the corresponding unstimulated control condition.

**Table S2**

| Experiment | $V_{½act}$, mV | $K_{act}$, mV | $V_{½SSI}$, mV | $K_{SSI}$, mV | $I_{Na,L}$, % | Apparent $\tau_{fast}$, ms |
|---|---|---|---|---|---|---|
| **Untransfected NRVMs** | | | | | | |
| Control | -40.6 ± 2.5 | 9.8 ± 0.9 | -70.8 ± 2.5 | 11.2 ± 1.6 | 0.46 ± 0.52 | 12.0 ± 1.2 |
| diC8-$PIP_2$ | -42.8 ± 2.1 | 9.6 ± 1.1 | -72.1 ± 2.7 | 12.0 ± 1.5 | 0.31 ± 0.47 | 10.8 ± 1.3 |
| **CRY2-PJ NRVMs** | | | | | | |
| Control | -40.2 ± 2.3 | 9.6 ± 0.8 | -69.2 ± 3.4 | 10.3 ± 1.4 | 0.48 ± 0.59 | 11.9 ± 1.0 |
| Control + blue light | -50.1 ± 2.5** | 10.8 ± 1.1 | -76.8 ± 2.8** | 11.6 ± 1.4 | 3.3 ± 0.75*** | 15.2 ± 1.1** |
| diC8-$PIP_2$ | -38.2 ± 2.4 | 9.4 ± 1.0 | -72.0 ± 2.6 | 12.0 ± 2.4 | 0.36 ± 0.62 | 10.3 ± 1.2 |
| diC8-$PIP_2$ + blue light | -38.5 ± 2.6 | 9.2 ± 0.9 | -69.7 ± 2.0 | 11.3 ± 1.8 | 0.34 ± 0.58 | 10.5 ± 1.6 |

**Table S2. $PIP_2$-dependent regulation of $Na_V$ current gating and late sodium current in neonatal rat ventricular myocytes.** Whole-cell sodium currents were recorded from primary neonatal rat ventricular myocytes under the indicated experimental conditions. Conductance-voltage relationships were generated from peak sodium current amplitudes and fitted with a Boltzmann function to determine the voltage of half-maximal activation, V½act, and slope factor, kact. Steady-state inactivation relationships were generated using the voltage-clamp protocols described in the Methods and fitted with a Boltzmann function to determine the voltage of half-maximal inactivation, V½SSI, and slope factor, kSSI. Untransfected control cells were recorded with or without 50 µM diC8-$PIP_2$ included in the pipette solution. CRY2-pseudojanin/CIBN-CAAX-transfected cells were recorded with or without 50 µM diC8-$PIP_2$ in the pipette solution and analyzed before and after 460 nm photoactivation. Gating data were obtained from 16 cells per untransfected control group and 12–16 cells per CRY2-pseudojanin group, collected across three independent biological replicates. $I_{Na,L}$ and apparent $\tau_{fast}$ were measured in separate cells. $I_{Na,L}$ was calculated as the mean current amplitude between 50 and 100 ms after peak current at −30 mV and is reported as a percentage of peak $I_{Na}$. Apparent $\tau f_{ast}$ was obtained from the same late-current recordings by fitting the decay phase of the current from the peak to 100 ms. $I_{Na,L}$ and apparent $\tau_{fast}$ data were obtained from 15–18 cells per group across three biological replicates. Values are means ± SD. Statistical significance was assessed using a t test with Welch's correction where ** is P < 0.01 and *** is P < 0.001.

77 **Table S3**

| Experiment | Peak $I_{Na}$ pA/pF | $I_{Na,L}$ pA/pF | $I_{Na,L}$, % of peak | $\tau_{fast}$, ms |
|---|---|---|---|---|
| **Untransfected iPS-CMs** | | | | |
| Control | −255.2 ± 46.2 | −0.89 ± 0.66 | 0.35 ± 0.25 | 6.6 ± 0.9 |
| Control + candesartan | −235.1 ± 44.4 | −0.96 ± 0.73 | 0.41 ± 0.30 | 6.8 ± 1.0 |
| Control + U73122 | −260.5 ± 53.9 | −0.96 ± 0.61 | 0.37 ± 0.22 | 6.7 ± 1.1 |
| Control + U73433 | −269.1 ± 49.4 | −0.67 ± 0.76 | 0.25 ± 0.28 | 6.6 ± 1.2 |
| Control + Staurosporine | −243.2 ± 44.1 | −0.73 ± 0.60 | 0.30 ± 0.24 | 6.5 ± 1.2 |
| Control + BAPTA | −232.0 ± 46.4 | −1.16 ± 0.67 | 0.50 ± 0.27 | 6.9 ± 1.1 |
| Control + diC8-$PIP_2$ | −221.5 ± 41.6 | −0.62 ± 0.35 | 0.28 ± 0.15 | 6.8 ± 0.5 |
| Ang II | −255.7 ± 51.6 | −8.08 ± 2.37*** | 3.16 ± 0.67*** | 10.2 ± 1.1** |
| Ang II + candesartan | −232.1 ± 47.2 | −1.00 ± 0.95 | 0.43 ± 0.40 | 6.9 ± 1.2 |
| Ang II + U73122 | −264.5 ± 47.7 | −1.06 ± 0.66 | 0.40 ± 0.24 | 6.6 ± 1.0 |
| Ang II + U73433 | −264.3 ± 55.0 | −7.88 ± 2.57*** | 2.98 ± 0.75*** | 11.1 ± 1.7** |
| Ang II + Staurosporine | −238.7 ± 44.5 | −7.30 ± 2.27*** | 3.06 ± 0.76*** | 10.6 ± 1.6** |
| Ang II + BAPTA | −272.6 ± 52.7 | −8.48 ± 2.39*** | 3.11 ± 0.64*** | 11.3 ± 1.6** |
| Ang II + diC8-$PIP_2$ | −225.1 ± 41.4 | −1.01 ± 0.51 | 0.45 ± 0.21 | 6.8 ± 0.6 |
| **M3q-DREADD iPS-CMs** | | | | |
| Control | −266.6 ± 54.4 | −0.59 ± 0.55 | 0.22 ± 0.20 | 6.2 ± 1.3 |
| Control + candesartan | −264.4 ± 55.3 | −0.82 ± 0.81 | 0.31 ± 0.30 | 6.8 ± 1.2 |
| Control + U73122 | −249.5 ± 54.6 | −0.67 ± 0.57 | 0.27 ± 0.22 | 6.9 ± 1.2 |
| Control + U73433 | −240.8 ± 48.7 | −0.77 ± 0.62 | 0.32 ± 0.25 | 6.5 ± 1.1 |
| Control + Staurosporine | −265.6 ± 54.4 | −0.77 ± 0.66 | 0.29 ± 0.24 | 7.0 ± 0.9 |
| Control + BAPTA | −267.4 ± 54.3 | −0.86 ± 0.74 | 0.32 ± 0.27 | 7.3 ± 1.4 |
| Control + diC8-$PIP_2$ | −258.8 ± 47.1 | −0.78 ± 0.44 | 0.30 ± 0.16 | 5.9 ± 1.3 |
| CNO | −232.5 ± 44.5 | −7.51 ± 1.94*** | 3.23 ± 0.56*** | 11.2 ± 1.3** |
| CNO + candesartan | −224.4 ± 42.5 | −7.14 ± 1.63*** | 3.18 ± 0.41*** | 11.3 ± 1.3** |
| CNO + U73122 | −225.6 ± 43.1 | −0.68 ± 0.56 | 0.30 ±0.24 | 7.2 ± 1.2 |
| CNO + U73433 | −255.0 ± 49.6 | −8.47 ± 2.25*** | 3.32 ± 0.60*** | 11.3 ± 1.1** |
| CNO + Staurosporine | −240.4 ± 45.3 | −7.67 ± 1.71*** | 3.19 ± 0.38*** | 10.9 ± 1.3** |

| | | | | |
|---|---|---|---|---|
| CNO + BAPTA | −234.7 ± 51.0 | −7.28 ± 1.96*** | 3.10 ± 0.49*** | 11.1 ± 1.2** |
| CNO + diC8-$PIP_2$ | −255.6 ± 52.2 | −0.89 ± 0.64 | 0.35 ± 0.24 | 6.7 ± 0.8 |
| **CRY2-PJ iPS-CMs** | | | | |
| Control, dark | −229.4 ± 48.0 | −0.87 ± 0.69 | 0.38 ± 0.29 | 6.9 ± 1.1 |
| Control + 460 nm | −229.0 ± 44.7 | −7.08 ± 1.90*** | 3.09 ± 0.57*** | 12.2 ± 1.4** |
| diC8-$PIP_2$, dark | −274.4 ± 56.4 | −0.71 ± 0.59 | 0.26 ± 0.21 | 6.7 ± 1.0 |
| diC8-$PIP_2$ + 460 nm | −250.6 ± 52.0 | −0.78 ± 0.50 | 0.31 ± 0.19 | 6.9 ± 1.2 |

**Table S3. Peak and late $I_{Na}$ parameters in human iPS-derived cardiomyocytes.** Whole-cell sodium currents were recorded from human iPS-derived cardiomyocytes under the indicated experimental conditions using the voltage-clamp protocols described in the Methods. Where indicated, cells were transfected to express the CNO-activated M3q-DREADD or light-activated CRY2-pseudojanin together with CIBN-CAAX. Peak $I_{Na}$ was measured during depolarization to −30 mV and expressed as current density. $I_{Na,L}$ was measured from the same traces as the mean current amplitude between 50 and 100 ms after the peak current at −30 mV. $I_{Na,L}$ is reported both as current density and as a percentage of peak $I_{Na}$, calculated as ($I_{Na,L}$/$I_{Na}$,peak) × 100. The apparent time constant of fast inactivation, τfast, was obtained by fitting the decay phase of the current from the peak to 100 ms with a single-exponential function with a non-zero plateau. Values are means ± SD from 12–22 cells per condition, collected from at least 3 independent biological replicates. The mean whole-cell capacitance value was 28 ± 5 pF. For direct comparisons in which currents were recorded from the same cell under two conditions, statistical significance was assessed using paired two-tailed t tests. For comparisons among multiple independent treatment groups, statistical tests are indicated in the table footnotes. Significance is indicated as **P<0.01, ***p < 0.001 versus the indicated control or stimulated condition.

**Table S4**

| Experiment | V½$_{act}$, mV | K$_{act}$, mV | V½$_{SSI}$, mV | K$_{SSI}$, mV |
|---|---|---|---|---|
| **Na$_V$1.5 variant with control or diC8-PIP$_2$ pipette** | | | | |
| Na$_V$1.5 | -45.2 ± 2.6 | 5.8 ± 0.7 | -81.4 ± 2.6 | 7.6 ± 0.5 |
| Na$_V$1.5 + diC8-PIP$_2$ (50 µM) | -44.8 ± 3.1 | 6.0 ± 1.0 | -80.0 ± 3.1 | 7.8 ± 0.7 |
| Na$_V$1.5 + diC8-PIP$_2$ (100 µM) | -42.5 ± 3.3$^{*}$ | 6.1 ± 1.1 | -78.1 ± 2.9 | 7.4 ± 1.1 |
| R1644C | -54.6 ± 4.5 | 5.1 ± 0.8 | -70.3 ± 3.5 | 6.4 ± 1.3 |
| R1644C + diC8-PIP$_2$ | -48.5 ± 4.3$^{***}$ | 5.8 ± 1.5 | -76.7 ± 3.3$^{**}$ | 6.8 ± 1.0 |
| R1644A | -55.1 ± 3.8 | 5.4 ± 1.2 | -69.6 ± 4.1 | 5.9 ± 1.1 |
| R1644A + diC8-PIP$_2$ | -49.6 ± 3.6$^{****}$ | 5.9 ± 2.1 | -77.2 ± 3.8$^{***}$ | 6.0 ± 1.2 |
| R1512W | -47.5 ± 5.0 | 5.2 ± 1.0 | -86.2 ± 3.1 | 6.5 ± 0.8 |
| R1512W + diC8-PIP$_2$ | -49.4 ± 6.2 | 6.3 ± 1.2 | -82.2 ± 3.3$^{**}$ | 6.7 ± 0.9 |
| **Na$_V$1.5 + CRY2-PJ with control or diC8-PIP$_2$ pipette** | | | | |
| Na$_V$1.5 control, dark | -45.5 ± 3.7 | 6.0 ± 0.8 | -80.4 ± 4.0 | 7.3 ± 0.9 |
| Na$_V$1.5 + blue light | -54.4 ± 2.6$^{****}$ | 9.3 ± 1.1 | -91.2 ± 6.3$^{****}$ | 11.2 ± 1.2$^{***}$ |
| Na$_V$1.5 + diC8-PIP$_2$, dark | -44.7 ± 3.5 | 6.2 ± 1.1 | -78.6 ± 3.7 | 6.9 ± 0.7 |
| Na$_V$1.5 + diC8-PIP$_2$ + blue light | -46.0 ± 3.2 | 6.6 ± 1.3 | -80.7 ± 4.2 | 7.1 ± 0.8 |
| R1644C, dark | -53.3 ± 3.7 | 6.4 ± 0.9 | -72.5 ± 4.1 | 6.6 ± 0.8 |
| R1644C + blue light | -59.9 ± 2.1$^{***}$ | 6.1 ± 1.0 | -74.0 ± 4.0$^{***}$ | 8.7 ± 1.7$^{**}$ |
| R1644C + diC8-PIP$_2$, dark | -48.2 ± 1.7 | 5.5 ± 0.5 | -76.0 ± 4.2 | 6.3 ±1.1 |
| R1644C + diC8-PIP$_2$ + blue light | -47.0 ± 4.2 | 5.8 ± 1.1 | -74.1 ± 4.0 | 6.8 ± 1.3 |
| **Na$_V$1.5 + M3q DREADD with control or diC8-PIP$_2$ pipette** | | | | |
| Na$_V$1.5 control | -47.0± 3.5 | 6.1 ± 0.9 | -80.4 ± 4.1 | 7.4 ± 1.0 |
| Na$_V$1.5 + CNO | -55.2 ± 3.9$^{****}$ | 8.7 ± 1.6$^{**}$ | -90.7 ± 3.5$^{****}$ | 10.8 ± 1.1$^{***}$ |
| Na$_V$1.5 + diC8-PIP$_2$ | -44.8 ± 4.2 | 5.8 ± 1.2 | -79.6 ± 4.4 | 7.2 ± 0.7 |
| Na$_V$1.5 + diC8-PIP$_2$ + CNO | -45.0 ± 3.5 | 6.0 ± 1.2 | -78.6 ± 3.7 | 7.3 ± 0.9 |
| R1644C | -55.6 ± 4.8 | 5.7 ± 1.3 | -74.3 ± 6.2 | 8.3 ± 1.1 |
| R1644C + CNO | -58.1 ± 3.3$^{**}$ | 6.3 ± 1.1 | -78.0 ± 6.8$^{**}$ | 7.7 ± 1.2 |
| R1644C + diC8-PIP$_2$ | -48.1 ± 2.0 | 6.0 ± 0.6 | -75.4 ± 3.9 | 6.9 ± 1.0 |
| R1644C + diC8-PIP$_2$ + CNO | -49.1 ± 2.8 | 6.9 ± 0.8 | -74.2 ± 4.5 | 6.7 ± 0.9 |

**Table S4. Voltage dependence of activation and steady-state fast inactivation of WT and mutant $Na_V1.5$ channels expressed in HEK293T cells.** Whole-cell sodium currents were recorded from HEK293T cells transiently expressing human $Na_V1.5$ WT, R1644C, R1644A, or R1512W channels, as indicated. Cells were co-transfected with $Na_V\beta1$ and iRFP in all experiments, allowing identification of transfected cells without photoactivation of the optogenetic machinery. Where indicated, the water-soluble $PIP_2$ analog diC8-$PIP_2$ was included in the patch pipette solution. For CRY2-PJ experiments, cells were additionally co-transfected with the optogenetic $PIP_2$ phosphatase CRY2-PJ, and currents were recorded before and after illumination with blue light, with or without diC8-$PIP_2$ in the pipette solution. Peak sodium currents were measured during depolarizing voltage steps and converted to conductance using the calculated sodium reversal potential. Conductance-voltage relationships were normalized to maximal conductance and fit with a Boltzmann function to determine the voltage of half-maximal activation, V½act, and the activation slope factor, kact. Steady-state fast inactivation was determined using a standard conditioning pulse protocol followed by a test pulse to measure channel availability. Normalized availability curves were fit with a Boltzmann function to determine the voltage of half-maximal steady-state inactivation, V½SSI, and the inactivation slope factor, kSSI. Data are presented as mean ± SD from 15-28 cells per condition from 3-6 independent transfections. Statistical comparisons were performed using paired t tests for within-cell comparisons before and after blue light illumination, and unpaired t tests with Welch's correction for comparisons between independent pipette or mutant conditions, as appropriate. *$P < 0.05$, **$P < 0.01$, ***$P < 0.001$, and ****$P < 0.0001$ versus the corresponding control condition.

**Table S5**

| Experiment | Peak $I_{Na}$ pA/pF | $I_{Na,L}$ pA/pF | $I_{Na,L}$, % of peak | $\tau_{fast}$, ms |
|---|---|---|---|---|
| **Na$_V$1.5 variant with control or diC8-PIP$_2$ pipette** | | | | |
| Na$_V$1.5 | −323.5 ± 22 | −1.1 ± 0.5 | 0.35 ± 0.15 | 6.6 ± 0.9 |
| Na$_V$1.5 + diC8-PIP$_2$ (50 μM) | −338.2 ± 18 | −1.3 ± 0.5 | 0.37 ± 0.14 | 6.8 ± 1.0 |
| Na$_V$1.5 + diC8-PIP$_2$ (100 μM) | −332.5 ± 20 | −1.0 ± 0.3 | 0.30 ± 0.10 | 6.7 ± 1.1 |
| R1644C | −147.8 ± 37 | −4.0 ± 1.0 | 2.70 ± 0.20 | 5.6 ± 1.0 |
| R1644C + diC8-PIP$_2$ | −158 ± 41 | −1.7 ± 0.6 | 1.10 ± 0.21**** | 5.4 ± 1.3 |
| R1644A | −275.5 ± 53 | −8.8 ± 2.1 | 3.20 ± 0.46 | 5.5 ± 1.2 |
| R1644A + diC8-PIP$_2$ | −255 ± 42 | −3.4 ± 1.0 | 1.35 ± 0.32**** | 5.8 ± 1.2 |
| K1641A | −26.8 ± 11 | N.D. | N.D. | N.D. |
| K1641A + diC8-PIP$_2$ | −24.3 ± 4 | N.D. | N.D. | N.D. |
| R1512W | −137.5 ± 17 | −0.8 ± 0.3 | 0.55 ± 0.20 | 7.4 ± 1.6 |
| R1512W + diC8-PIP$_2$ | −123.6 ± 22 | −0.6 ± 0.2 | 0.45 ± 0.12 | 7.7 ± 1.4 |
| **Na$_V$1.5 + CRY2-PJ with control or diC8-PIP$_2$ pipette** | | | | |
| Na$_V$1.5 control, dark | −312 ± 18 | −1.3 ± 0.7 | 0.42 ± 0.23 | 7.0 ± 0.8 |
| Na$_V$1.5 + 460 nm | −326 ± 32 | −11.1 ± 1.7 | 3.40 ± 0.41**** | 12.4 ± 1.8**** |
| Na$_V$1.5 + diC8-PIP$_2$, dark | −352 ± 28 | −1.4 ± 0.9 | 0.39 ± 0.25 | 6.7 ± 0.9 |
| Na$_V$1.5 + diC8-PIP$_2$ + 460 nm | −347 ± 25 | −1.4 ± 1.1 | 0.41 ± 0.32 | 7.1 ± 1.4 |
| R1644C, dark | −139 ± 20 | −4.0 ± 0.8 | 2.90 ± 0.36 | 6.1 ± 0.6 |
| R1644C + 460 nm | −131 ± 33 | −5.5 ± 1.5 | 4.20 ± 0.44*** | 8.4 ± 1.2** |
| R1644C + diC8-PIP$_2$, dark | −142 ± 37 | −4.4 ± 1.3 | 3.10 ± 0.39 | 5.8 ± 0.7 |
| R1644C + diC8-PIP$_2$ + 460 nm | −146 ± 34 | −5.3 ± 1.4 | 3.60 ± 0.47* | 7.9 ± 1.4* |

**Table S5. Late sodium current and fast-inactivation kinetics of WT and mutant Na$_V$1.5 channels expressed in HEK293T cells.** Whole-cell sodium currents were recorded from HEK293T cells transiently expressing human Na$_V$1.5 WT, R1644C, R1644A, K1641A, or R1512W channels, as indicated. Cells were co-transfected with Na$_V$β1 and iRFP in all experiments, allowing identification of transfected cells without photoactivation of the optogenetic machinery. Currents were elicited by depolarizing voltage steps to −30 mV from a negative holding potential. Where indicated, the water-soluble PIP$_2$ analog diC8-PIP$_2$ was included in the patch pipette solution. For CRY2-PJ experiments, cells were additionally co-transfected with the optogenetic PIP$_2$-phosphatase CRY2-PJ, and currents were recorded before and after illumination with 460 nm blue light, with or without diC8-$PIP_2$ in the pipette solution. Peak $I_{Na}$ density was calculated from the maximal inward current normalized to cell capacitance. Late sodium current density ($I_{Na,L}$) was measured as the mean current between 50 and 100 ms after the peak current and normalized to cell capacitance. $I_{Na,L}$ is also expressed as a percentage of the corresponding peak current measured in the same cell, calculated using absolute current amplitudes. The fast component of macroscopic inactivation was quantified by fitting the decay phase of the current from the peak to 100 ms with a single-exponential function plus a non-zero steady-state component. N.D., not determined because current amplitudes were too small for reliable measurement of late current or decay kinetics. Data are presented as mean ± SD from 12–45 cells per condition from 3–5 independent transfections. Statistical comparisons were performed using paired t tests for within-cell comparisons before and after 460 nm illumination, and unpaired t tests for comparisons between independent pipette or mutant conditions, as appropriate. *$P < 0.05$, **$P < 0.01$, ***$P < 0.001$, and ****$P < 0.0001$ versus the corresponding control condition.

**Figure S1**

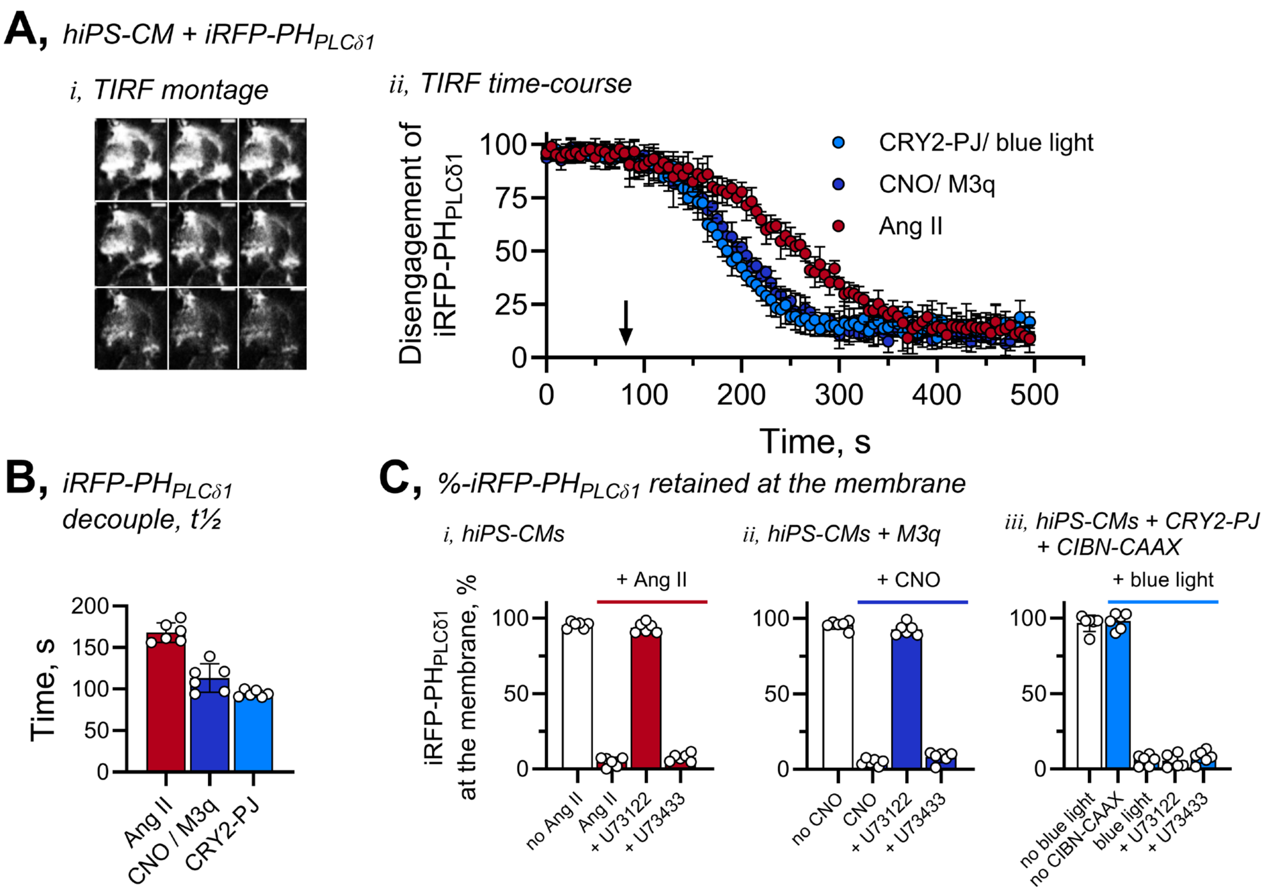


**Figure S1. Total internal reflection fluorescence (TIRF) analysis of iRFP-PH$_{PLC\delta1}$ membrane decoupling in iPS-CMs.** Representative TIRF image series and summary time courses show loss of plasma-membrane iRFP-PH$_{PLC\delta1}$ fluorescence after Ang II stimulation, M3q/CNO activation, or CRY2-PJ photoactivation. U73122 prevents receptor-driven biosensor decoupling, whereas U73433 does not. Control experiments include unstimulated cells and optogenetic-component controls as indicated. Data are mean ± SD from the 6 – 8 cells per condition studies in three independent experiments.

**Figure S2.**

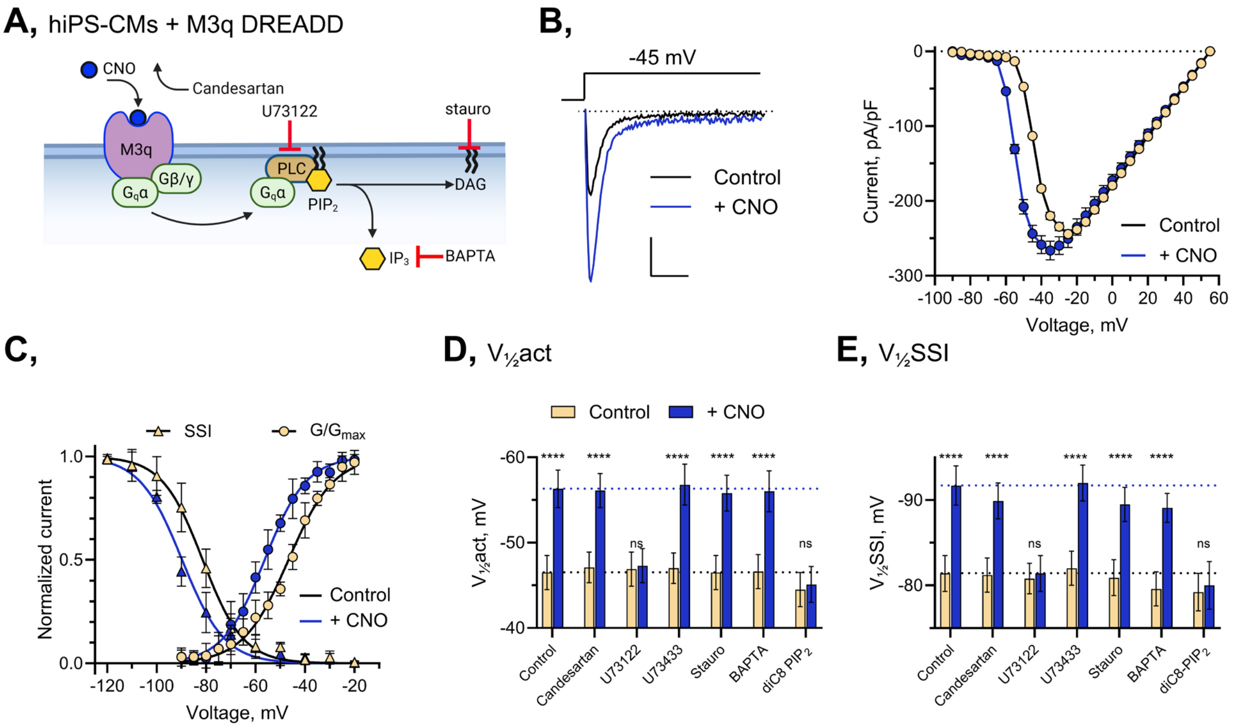


**Figure S2. Gq-PLC signaling via the M3q DREADD shifts Nav1.5 gating through $PIP_2$ depletion.** **(A)** Schematic of the M3q-DREADD/CNO strategy. **(B)** Representative iPS-CM $I_{Na}$ traces before and after CNO. **(C)** Current-voltage, conductance-voltage, and steady-state inactivation relationships show CNO-induced hyperpolarizing shifts in activation and availability. **(D, E)** Summary analyses show that U73122 and intracellular diC8-$PIP_2$ prevent the CNO response, whereas candesartan, U73433, staurosporine, and BAPTA do not. Data are mean ± SD; n values and statistical comparisons are reported in Tables S1 and S3.

**Figure S3**

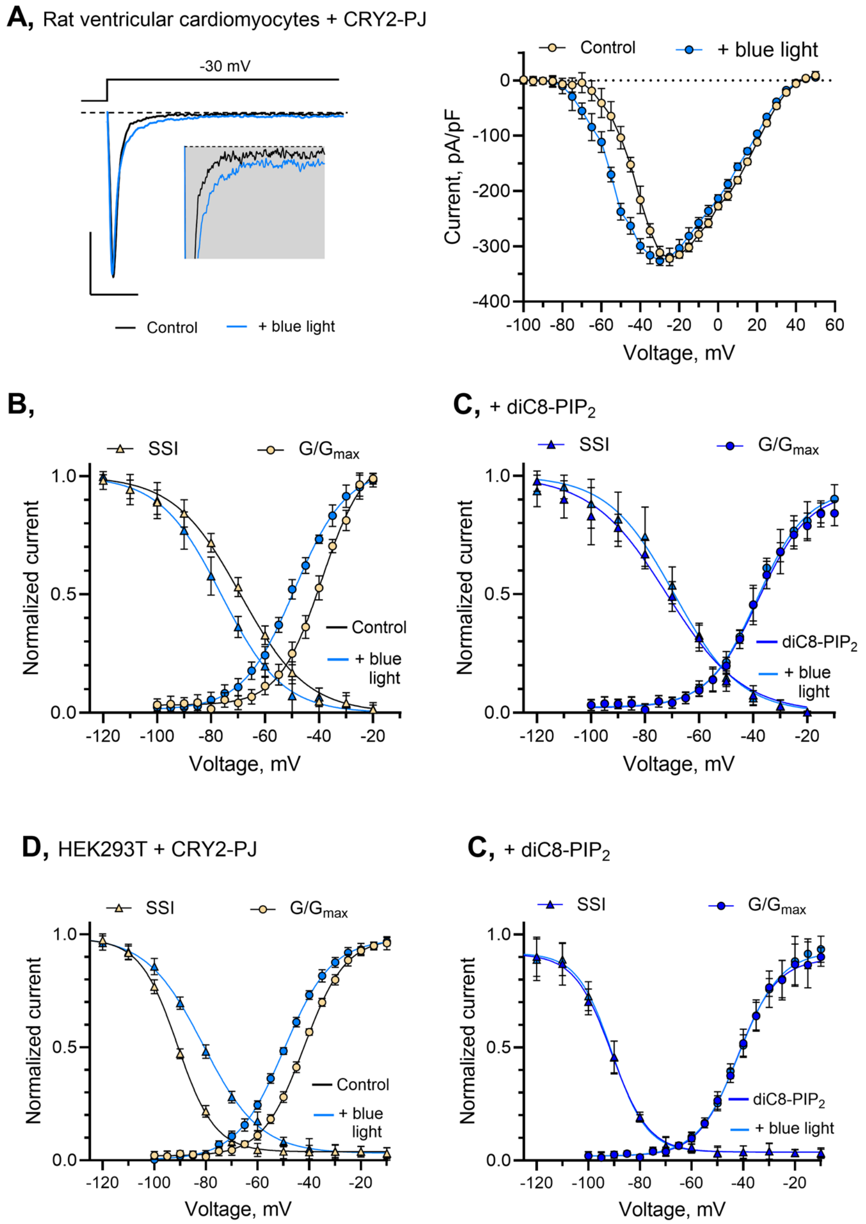


**Figure S3. Acute $PIP_2$ depletion shifts $I_{Na}$ activation and inactivation in neonatal rat ventricular cardiomyocytes and HEK293T cells. (A-C)** Representative currents and summary gating analyses show that CRY2-PJ photoactivation produces coordinated hyperpolarizing shifts in $V_{½}$act and $V_{½}$SSI in neonatal rat ventricular myocytes, and that 50 µM intracellular diC8-$PIP_2$ prevents these effects. Companion HEK293T experiments show a similar CRY2-PJ-dependent $Na_V1.5$ gating phenotype in a heterologous system. Data are mean ± SD; n values and statistical comparisons are reported in Tables S2 and S4.

**Figure S4**

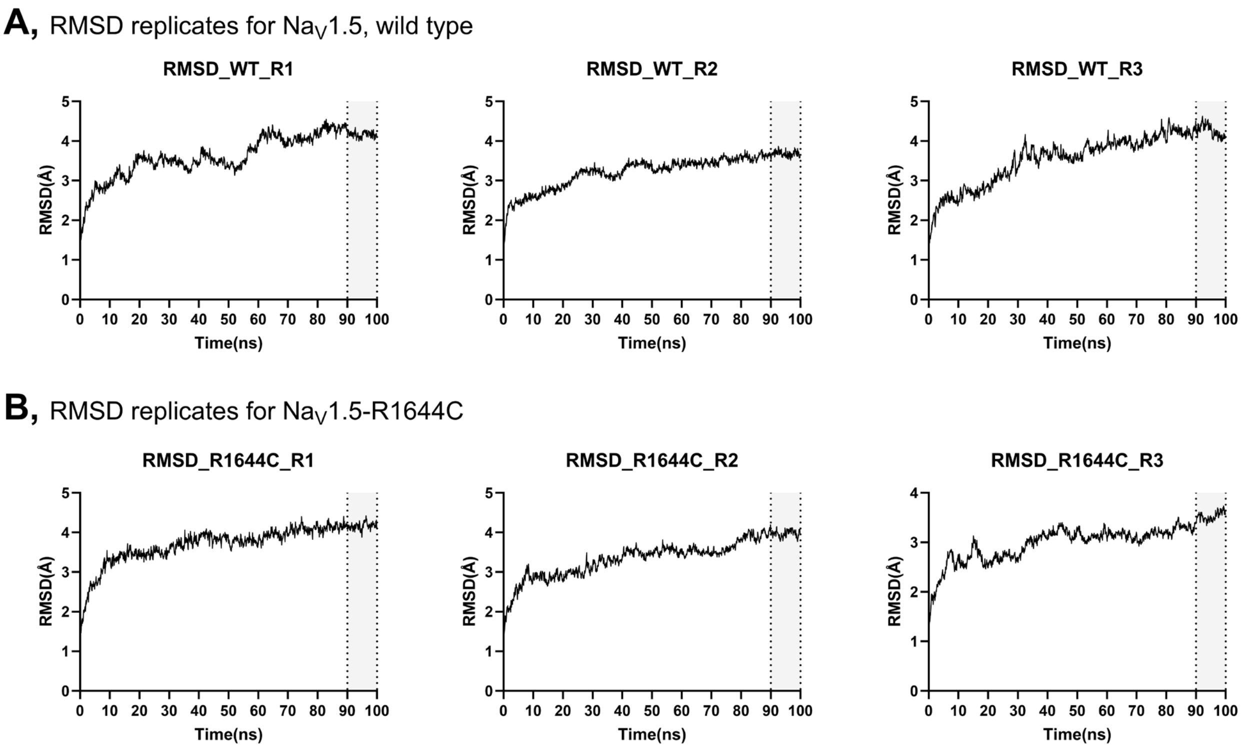


**Figure S4. Root-mean-square deviation (RMSD) validation of $Na_V1.5$-$PIP_2$ molecular dynamics simulations**. RMSD trajectories are shown for **(A)** three independent wild-type $Na_V1.5$-$PIP_2$ simulations and **(B)** three independent $Na_V1.5$-R1644C-$PIP_2$ simulations. These analyses support equilibration and trajectory stability over the production simulations used for contact and interaction-energy analyses.

### Figure S5

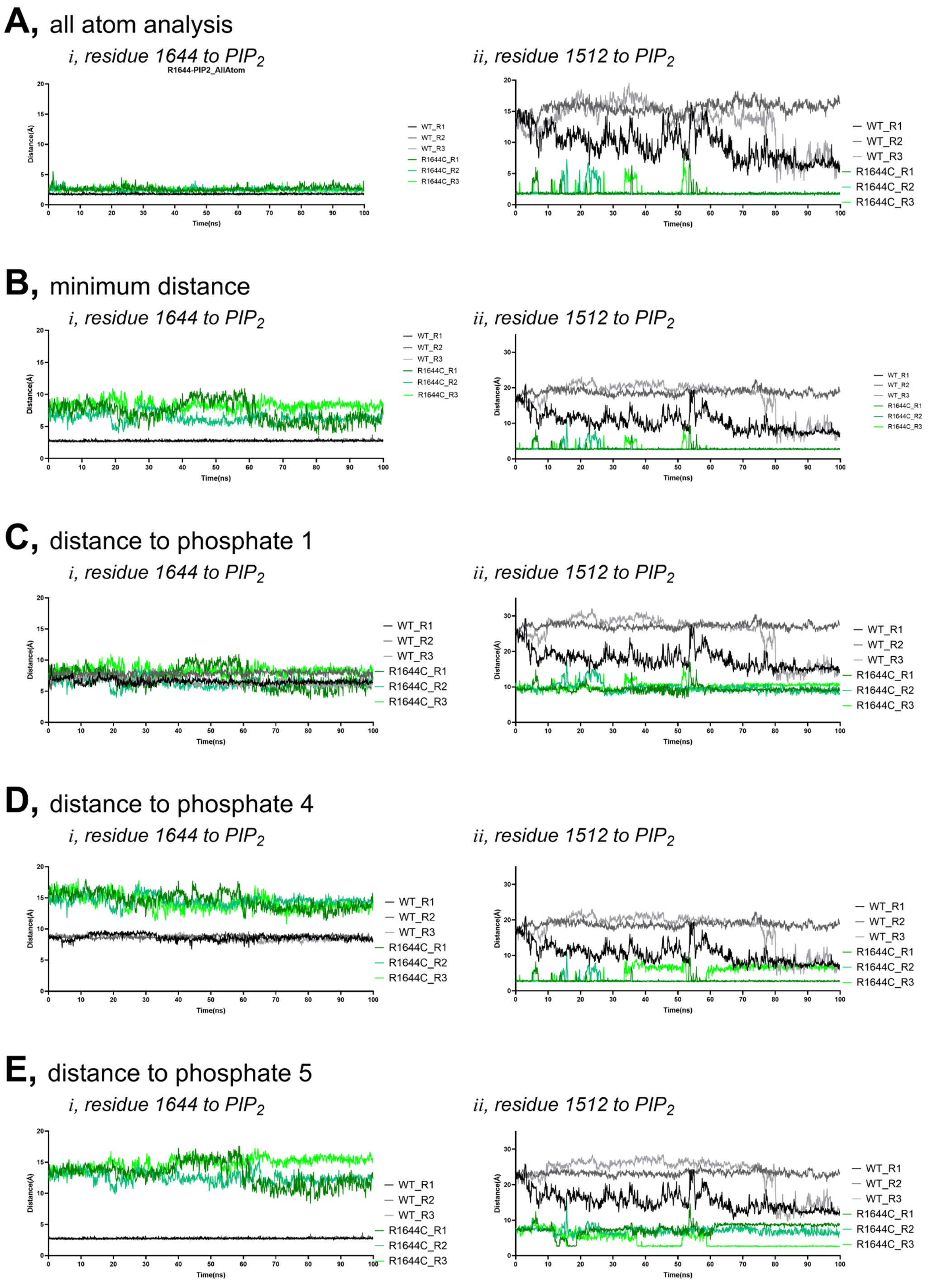


**Figure S5. Replicate molecular dynamics trajectories supporting $PIP_2$-contact analysis.** Replicate trajectories show minimum distances and atom contacts between $PIP_2$ and key residues in the wild-type and R1644C systems, including R1644, R1512, and phosphate-specific contacts. Analyses support loss of R1644-centered contacts and redistribution toward R1512 in the R1644C model.
